# Learning induces activation-mechanism–dependent neural plasticity in an intracortical microstimulation task

**DOI:** 10.64898/2026.06.05.730421

**Authors:** Robin Kim, Roy Lycke, Pavlo Zolotavin, Jon Montes, Chong Xie, Lan Luan

**Affiliations:** Department of Electrical and Computer Engineering, Rice University; Rice Neuroengineering Initiative, Rice University; Department of Bioengineering, Rice University

## Abstract

Electrical microstimulation provides high-resolution control of neural circuits for causal studies and restoration of impaired functions, yet how responses to artificial activation evolve with learning remains unclear. Here, we deploy a detection task and pair ultraflexible electrodes for stable intracortical microstimulation (ICMS) with longitudinal imaging and recordings to track single-cell and population responses across weeks of learning. Detection thresholds decreased with learning, indicating plasticity. Chronic imaging showed that stimulus-evoked recruitment expanded at a fixed current, while a consistent number of neurons continued to underlie behavioral responses. A subset of learning-sensitive cells enhanced modulation and reduced latency. Electrophysiological recordings further distinguished two forms of adaptation: directly activated, pulse-locked neurons strengthened their excitability, whereas polysynaptically recruited neurons expanded in number and were predictive of behavioral outcomes. These results show that learning in an ICMS task reshapes cortical circuits through activation-mechanism–dependent plasticity, underscoring the need for stimulation paradigms that adapt to both cell-intrinsic and network dynamics.

**One-sentence summary:** Learning alters neural responses to ICMS, revealing differences between direct activation and network activity linked to behavior.

## Introduction

The ability to alter neural activity and understand its effects is essential for understanding and augmenting brain function. Electrical microstimulation using penetrating electrodes provides a direct means of manipulating local activity and has been applied across diverse contexts, from mapping causal relationships in basic neuroscience to restoring sensory functions through brain–machine interfaces(*1*). In both scientific and clinical domains, the effects of microstimulation often unfold within the context of learning(*2–4*), which determines how artificial inputs are interpreted, incorporated, and ultimately stabilized within neural circuits.

A large body of work has characterized how intracortical microstimulation (ICMS) activates local and downstream neurons using optical imaging(*5*), electrophysiological recording(*6*), and functional MRI(*7*) across animal models and in human patients(*8*, *9*). These studies have illuminated key aspects of the activation process, such as spatially distributed direct activation(*5*) and temporally diverse responses arising from direct or synaptic recruitment(*9*, *10*). Yet, because these observations mostly come from single-session experiments conducted without behavioral learning(*10–13*), it remains unclear how ICMS-evoked activity evolves with time or how learning shapes the underlying circuit dynamics.

Learning is a powerful driver of neural plasticity, capable of modifying intrinsic excitability, synaptic strengths, and population representations to support new behaviors(*14*, *15*). Previous studies on the neural substrate of learning artificial inputs have primarily relied on optogenetic stimulation combined with Ca²⁺ imaging readouts(*16*, *17*). These studies showed that pairing artificial activation with behavioral training can amplify cortical responses, with neurons exhibiting progressively larger and more robust activity patterns as animals learn to interpret novel stimulation(*16*). In parallel, learning has been associated with increased stability of population representations across trials and sessions, indicating that artificial inputs can be integrated into consistent and reliable cortical activity patterns over time(*17*).

Both animal and human studies of ICMS have shown that behavioral detection thresholds and perceptual sensitivity can evolve over repeated stimulation and training(*3*, *18–26*), suggesting the presence of learning-related plasticity(*24*). In particular, a long-term human study reported gradual improvements in perception and task performance with chronic ICMS(*3*), highlighting the clinical relevance of adaptation to artificial inputs. However, findings from optogenetic stimulations do not directly translate to electrical microstimulation. Unlike optogenetics, which relies on genetic modification and produces stereotyped activation in genetically defined populations, ICMS recruits neurons through the spread of an electrical field, leading to mixed activation of somata, dendrites, and axons of passage(*27*). While this enables direct clinical translation, it also raises fundamental questions about how learning interacts with ICMS: when behavioral detection thresholds change during training, does this reflect altered excitability of the same neurons, the recruitment of new populations, or shifts in the balance between direct and network-mediated responses (Fig. 1A)? These possibilities reflect distinct forms of circuit adaptation, yet their contributions during ICMS learning remain unknown. Clarifying how learning shapes ICMS-evoked responses is therefore critical not only for understanding basic mechanisms of plasticity but also for informing the development of stimulation-based therapies and sensory brain-machine interfaces, where long-term efficacy depends on the brain’s ability to adapt to and incorporate artificial inputs.

**Fig. 1.**
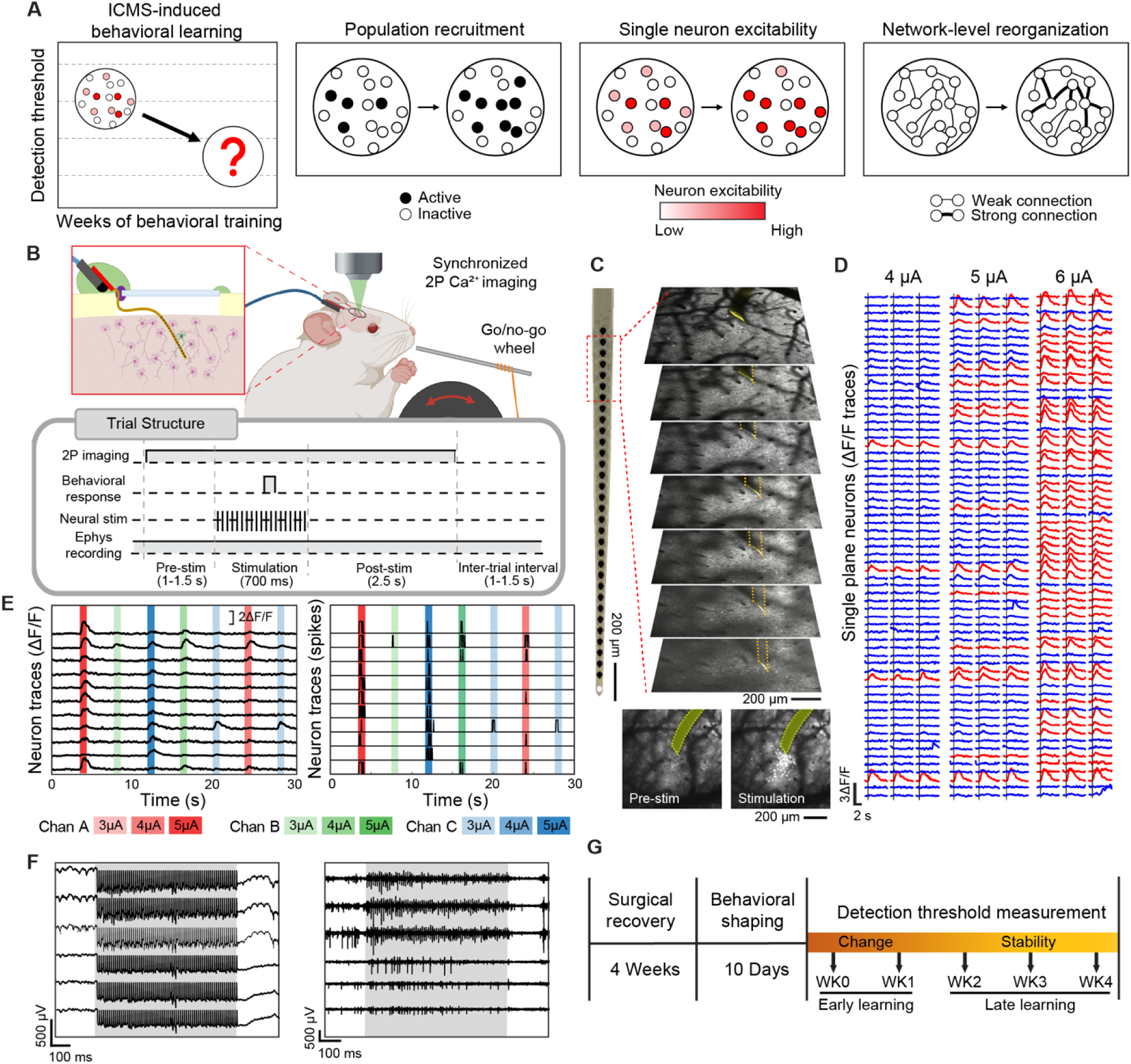
Concurrent imaging and electrophysiological recording reveal cellular activation and spiking dynamics during ICMS learning. (**A**) Behavioral detection thresholds evoked by ICMS change with training, indicating learning, and may arise from three mutually non-exclusive mechanisms of plasticity: (1) alterations in population recruitment, (2) changes in single-neuron excitability, and (3) network-level reorganization. (**B**) Schematics of multimodal, longitudinal measurements. Mice were implanted with a cranial window and an ultraflexible NET for both ICMS and recording (inset). During the ICMS detection task, mice reported perceived stimuli by turning a wheel while synchronized two-photon (2P) Ca²⁺ imaging and intracortical recordings were performed. Bottom diagram details the temporal alignment of all modalities within a single behavioral trial. Top schematic was created in BioRender. R. Lycke (2026); https://BioRender.com/gg4q9jv. (**C**) Representative volumetric Ca^2+^ imaging. Yellow dashed lines mark the NET at each imaging plane. Bottom: Maximum intensity projection of spontaneous activity (pre-stim) and neural activation at 4 μA. (**D**) Representative Ca^2+^ fluorescence from multiple cells in one imaging plane, with ICMS-evoked neurons in red and non-responding neurons in blue. **(E**) Representative Ca^2+^ transients (left) and deconvolved spikes (right) across multiple stimulation conditions. Colors denote stimulation channels; shading indicates current intensity. (**F**) Representative raw (left) and artifact-removed electrophysiological signals (right) demonstrating detection of spiking activity during stimulation at 100 Hz (gray-shaded period). (**G**) Experimental timeline, where early learning phases are expected to show significant changes in behavioral sensitivity, whereas late learning phases exhibit stability. Data from one representative animal for **C**-**F**.

Despite long-standing interest, chronic studies of ICMS-evoked activity at single-cell resolution and large-population scale during learning have been limited by technical barriers. Optical readouts in transgenic animals have been the mainstay for longitudinal population tracking(*28*), but they are poorly compatible with conventional stimulation electrodes over extended periods, restricting most stimulation-imaging experiments to acute sessions. Stable activation and measurement of the same neurons over time requires an exceptionally reliable tissue-electrode interface(*29*). Because stimulation often recruits neurons through axonal passage(*5*), its responses are highly sensitive to subtle changes at the interface, making it difficult to distinguish genuine circuit plasticity from interface drift. Furthermore, the two dominant single-cell readout methods, electrophysiology and optical imaging, each provide a partial view of evoked activity(*30*): optical imaging offers broad spatial coverage and reliable cell tracking but lacks temporal precision, whereas electrophysiology resolves millisecond dynamics but samples sparsely(*31*). These limitations have left learning-dependent evolution of ICMS-evoked neural circuit dynamics largely unexplored.

In this study, we address these limitations by establishing stable ICMS over chronic timescales and integrating it with multimodal measurements capable of following neuronal populations across weeks of behavioral learning. Using ultraflexible penetrating electrodes for ICMS and electrophysiological recording, integrated with two-photon calcium imaging, we captured stimulation-evoked activity with both single-cell spatial resolution and millisecond temporal precision. The ultraflexibility of the penetrating electrodes enables concurrent optical imaging(*32*) while maintaining a chronically stable tissue–electrode interface(*22*, *33*), minimizing variability in stimulation output and neural readout across sessions. This integrated approach enables direct linkage between behavioral learning and multiple measures of neural plasticity over time: two-photon calcium imaging quantifies population recruitment and spatial organization, while electrophysiological recordings resolve millisecond-scale spiking to assess neuronal excitability and infer activation mechanisms, together enabling dissection of single-neuron, population, and network-level adaptations. Using this platform, we find that ICMS-evoked activations are not fixed but undergo learning-dependent adaptation, with distinct forms of plasticity emerging across individual neurons and populations. These findings highlight that the integration of artificial inputs into neural circuits depends on both behavioral context and activation mechanisms, providing new insight into how stimulation drives adaptive plasticity and informing strategies for sensory brain-machine interfaces.

## Results

### Multimodal neural interface simultaneously detected cell-specific and spike-resolved activations during behavioral learning

During ICMS-induced learning, mice reliably detected microstimulation delivered to the primary somatosensory cortex (S1) and progressively improved their performance in a wheel-turning go/no-go task (Fig. 1B). Stimulation trains at 100 Hz elicited consistent behavioral responses within the 700-ms window, providing sufficient time for animals to respond within the stimulation window (fig. S1A, B). We employed ultraflexible nanoelectronic threads (NETs) for both stimulation and simultaneous spike recording, which offers two key technological advantages for studying the neural plasticity associated with ICMS learning(*22*). First, NETs are flexible and nearly transparent, enabling co-implantation with a cranial window and simultaneous volumetric two-photon imaging alongside electrophysiological recording (Fig. 1C shows the relative locations of NET and imaging volume). This multimodal configuration captures population activity at single-neuron resolution while resolving millisecond-scale spiking dynamics throughout the full cortical depth. Second, NETs maintain a stable tissue interface across various timescales, with minimal glial encapsulation over months(*22*, *34*) and within individual sessions, even in highly mobile tissue(*35*). This improved stability minimizes confounds from changes at the tissue-electrode interface, ensuring consistent stimulation output at fixed current levels and high-fidelity readout of evoked responses.

Within each animal, three NET contacts were designated as stimulation sites and used consistently across all sessions. During training, detection thresholds shifted both within and across sessions. To concentrate measurements around peri-threshold levels, we calculated the detection threshold every 100 trials (referred to as a block) within each session, and adjusted current amplitudes after each block to compensate for these in-session and across-session shifts in psychometric sensitivity (fig. S1C). Grouping trials by fixed current amplitude further showed systematic strengthening of evoked population responses with increasing currents (Fig. 1D). Randomization of stimulation contacts evoked distinct activity patterns (Fig. 1E), confirming the high selectivity of NET-evoked ICMS(*22*) and the fidelity of our optical readout.

Simultaneous electrophysiology and Ca²⁺ imaging during behavior provided complementary views of cortical dynamics (Fig. 1F). After artifact suppression, spike-resolved recordings revealed depth-dependent activation and spike timing precision that were not accessible from imaging alone, while optical recordings captured population activity at the single-cell resolution within the 1 × 1 × 0.4 mm³ field of view. Experiments began after a surgical recovery period(*36*) and ten days of behavioral shaping (Fig. 1G) and lasted for four weeks, a duration chosen based on our prior work(*22*) and those of others(*2*, *19*, *23*, *37*) showing that perceptual thresholds typically evolve and stabilize within this timescale.

### Behavioral learning enhances ICMS-evoked population activation over time

Over the weeks of behavioral learning, detection thresholds decreased (Fig. 2A and fig. S1D), with the steepest decline occurring during weeks 0–1, followed by more gradual changes and relative stability in weeks 2–4. Response times followed a similar time course (fig. S1E). Individual animals showed consistent behavioral trajectories with the group average (fig. S1D, E). Based on these behavioral dynamics, we refer to weeks 0–1 as the early learning phase and weeks 2–4 as the late learning phase. The detection thresholds in the late phase were significantly lower than those in the early phase (Fig. 2A), decreasing from a median of 5.5 µA to 4.2 µA (n_early_ = 43, n_late_ = 80, p = 7.68e-6, r = 0.40).

**Fig. 2.**
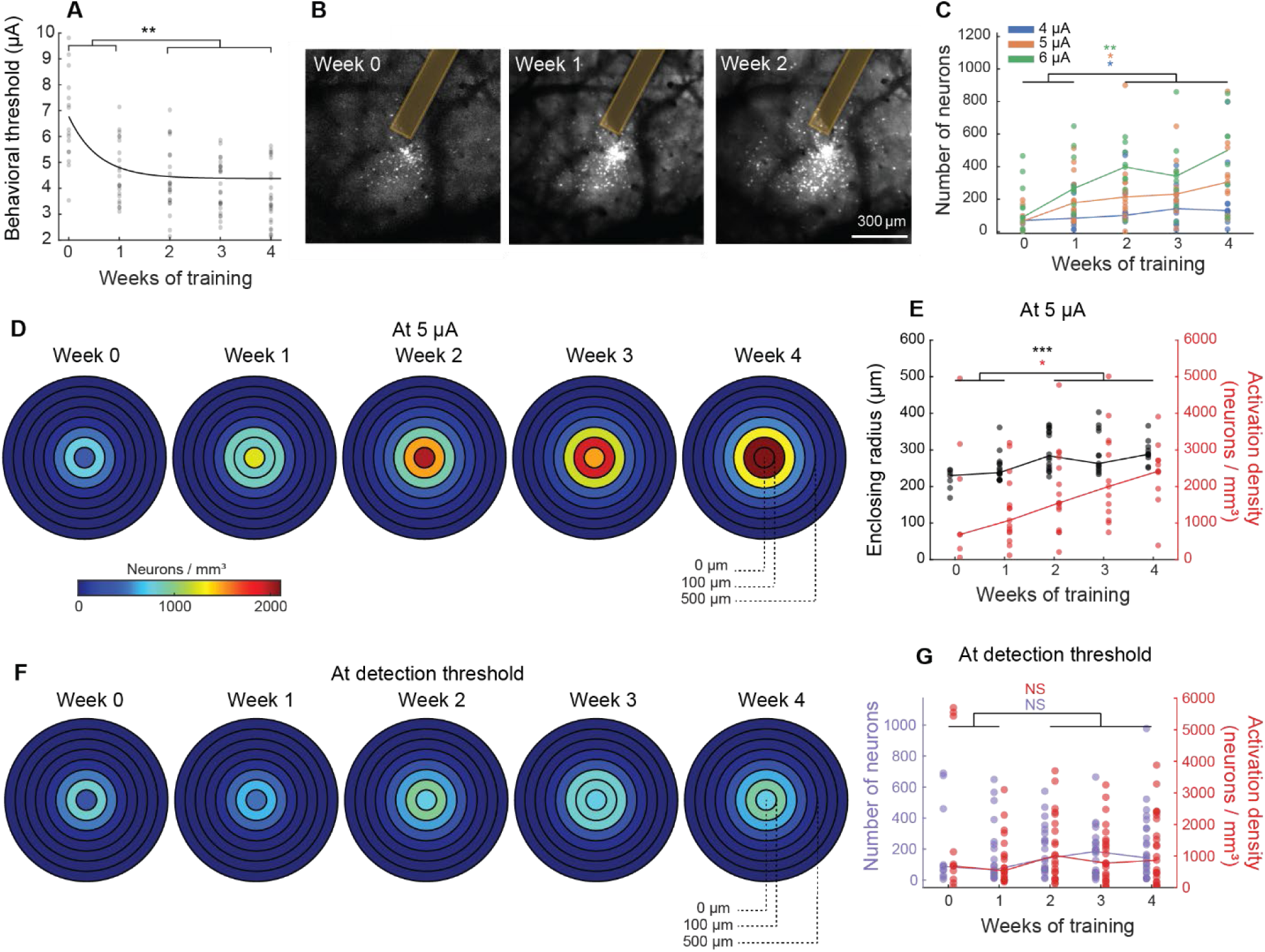
Behavioral learning enhances ICMS-evoked population activation over time. (**A**) Behavioral detection threshold decreased with learning. Dots represent individual stimulation channels; solid line shows an exponential fit. (**B**) Representative maximum intensity projection (MIP) of volumetric Ca^2+^ imaging at 4 μA over time. Yellow bands mark NET. (**C**) The number of ICMS-evoked neurons increases with training across all animals and sessions. Color indicates stimulation currents. (**D**) Average cell activation density as a function of distance from the stimulation sites at 5 μA, showing increasing activation with time. (**E**) Both the activation density within 200 μm of the stimulation site (red) and the radius enclosing 50% of the activated population (black) show significant increases over training. (**F**) Average cell activation density at detection thresholds shows relatively stable activation across training weeks when stimulation currents were adjusted to match thresholds. (**G**) At the detection threshold, the activation density within 200 μm of the stimulation site (red) and total number of activated neurons (purple) remained stable. Each dot represents an activation metric in a session at that channel’s detection threshold current. In (**C**), (**E**), (**G**), each dot represents an activation metric for a given channel, session, and animal; lines represent the median. n = 6 animals, 18 stimulation channels for all panels except **B**. Two-sided Mann-Whitney U tests, NS, not significant, * p<0.05; ** p<0.01, and ***p<0.001.

To assess how the ICMS-evoked neural activation evolves with learning, we quantified the activation population under two-photon volumetric Ca^2+^ imaging for 6 mice (Methods). In one representative animal at a fixed stimulation current of 4 µA, the number of ICMS-activated neurons increased over time within the same field of view (FOV), with progressively more cells recruited from week 0 (71 neurons) to week 1 (114 neurons) and to week 2 (220 neurons) (Fig. 2B). The peri-threshold currents were comparable across animals, and the increase in the number of activated cells is consistent across all animals, sessions, and stimulation currents, resulting in significantly larger activated populations at all current levels in the late learning phase than in the early phase (Fig. 2C, 4 µA: n_early_ = 13, n_late_ = 41, p = 0.043, r = 0.28; 5 µA: n_early_ = 20, n_late_ = 35, p = 0.015, r = 0.33; 6 µA: n_early_ = 22, n_late_ = 26, p = 0.0069, r = 0.39).

We next examined whether this increase reflected spatially broader recruitment. For each activated neuron, we calculated its three-dimensional (3D) distance to the stimulation site and determined the activation density within concentric 100 μm shells centered on the stimulating site (Methods). The activation density peaked at around 200 μm (fig. S2A, B), consistent with previous studies in the absence of behavioral learning(*5*). Most of the learning-related change in activation density also occurred in the close vicinity of the stimulation site (Fig. 2D, fig. S2C), resulting in significantly higher activation density with learning within a radius of 200 μm (Fig. 2E, n_early_ = 22, n_late_ = 40; p = 0.015, r = 0.31). To quantify the spatial spread of activation, we defined the distance enclosing 50% of all activated neurons as the *activation radius* (Methods), which also increased over time (Fig. 2E, p = 1.1e-5, r = 0.56). Importantly, in control animals not undergoing ICMS-driven behavioral learning, both the number and spatial distribution of activated neurons remained stable and resembled those in the initial training weeks (fig. S2E, F), confirming that the expansion of the activated population is learning dependent.

Given that over time activation increased at a fixed current while perceptual thresholds decreased, we next examined whether neural recruitment changed at the perceptual threshold over training. Notably, when the stimulation current was adjusted to the behavioral detection threshold for each session (Fig. 2F, G), both the activation density and number of activated neurons remained stable across learning phases (n_early_ = 32, n_late_ = 69; density: p = 0.511, r = 0.07; total number: p = 0.253, r = 0.11). Thus, despite declining detection thresholds, a consistent number of neurons underlie behavioral responses throughout the learning period.

### Longitudinal tracking of individual neurons reveals learning-sensitive subpopulations

The overall increase in the activated population during behavioral learning implies recruitment of new neurons at the same current. We next asked whether some neurons remained consistently activated throughout the training and how their responses evolved over time compared to those recruited later. To address this, we used PatchWarp(*38*) to align images across sessions, and longitudinally track stimulation-activated neurons across weeks (Methods). Representative tracking within the same imaging plane (Fig. 3A) shows that some neurons were repeatedly recruited over weeks, whereas others appeared only once, a pattern particularly common in later weeks. Among 10,608 tracked neurons in 3 animals, 4469 neurons were activated by ICMS. Among the ICMS-activated neurons, 114 were activated at all weeks, 370 neurons were activated in 4 out of 5 weeks, and 2063 were only activated in one session (Fig. 3B). This relatively small fraction of stably activated neurons is expected, given the tight tissue integration of NETs. First, minimal glial scarring around NETs lowers behavioral detection thresholds by several-fold compared with rigid microelectrodes(*22*), resulting in the recruitment of relatively few neurons. In addition, because ICMS often drives neurons antidromically via their axons, activation patterns are highly sensitive to even subcellular changes at the tissue–electrode interface, further limiting the number of neurons that remain stably activated over time. The persistence of a stably activated subset across weeks thus underscores the exceptional interface stability afforded by NETs.

**Fig. 3.**
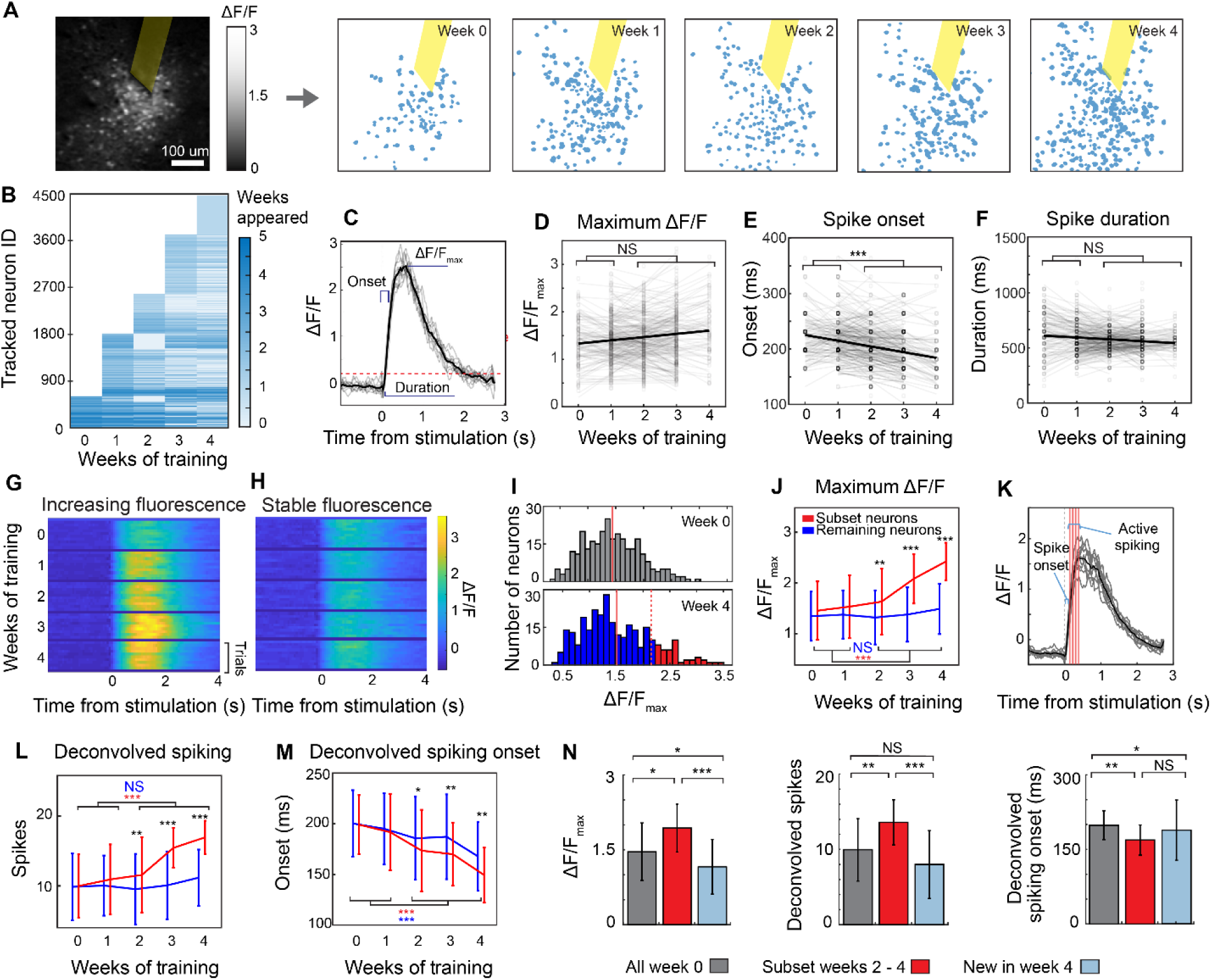
Longitudinal tracking of ICMS-evoked neurons reveals learning-sensitive subpopulations. (**A**) Representative tracking of ICMS-evoked neurons from single-plane imaging. Left: neural activation at 5 μA. Right: Binarized ROIs showing active neurons across weeks with NET as a yellow ribbon. (**B**) Number of neurons activated and tracked across all weeks. Neurons are sorted by their first appearance session and color-coded for tracking longevity. (**C**) Representative fluorescence traces of an ICMS-evoked neuron, denoting ΔF/F_max_, onset times, and response duration. The thick black line shows 10-trial average. (**D**-**F**) Longitudinal ΔF/F_max_ (**D**), onset times (**E**), and spiking duration (**F**) for tracked neurons. Solid lines show linear regression. (**G-H**) ΔF/F_max_ from two example neurons: one with increasing ΔF/F_max_ over training (**G**), and one remaining stable (**H**). (**I**) Distribution of ΔF/F_max_ of tracked neurons at weeks 0 and 4 from one representative animal. Solid and dashed vertical lines indicate the mean and mean + STD, respectively. Red marks the high-intensity subgroup; blue marks remaining neurons. (**J**) ΔF/F_max_as a function of time in training for the high-intensity subgroup (red) and remaining neurons (blue). **(K)** Example deconvolution of fluorescence traces for an ICMS-evoked neuron (thick black line: mean of 10 trials). Red vertical lines mark deconvolved spikes. (**L, M**) Deconvolved spike number **(L)** and onset time (**M**) as a function of time in training for the same two neuron groups shown in (**I**, **J**). (**N**) Comparison of ΔF/F_max_, deconvolved spike numbers and spike onset latency among the enhanced subgroup in the late phase (red), all longitudinally recruited neurons at week 0 (grey), and newly recruited neurons at week 4. n=3 animals for **D-F**, **J**, **L-N.** Wilcoxon signed-rank tests for **D-F**; two-sided Mann-Whitney U test for **J**, **L**, **M**; Kruskal-Wallis tests with Tukey-Kramer post-hoc correction for **N**. NS, not significant, * p<0.05; ** p<0.01, and *** p<0.001.

We next analyzed how the fluorescence response changed with learning, quantifying the maximum fluorescence intensity (ΔF/F_max_), onset time, and fluorescence duration (Fig. 3C) in each session. We first focused on neurons that were reliably activated at a given current and tracked across all sessions and all but one session in which the current was applied. On average, these longitudinally stable cells exhibited no significant changes in either ΔF/F_max_ or fluorescence duration (n = 226, Wilcoxon signed-rank tests; p = 0.8478, r < 0.01 for ΔF/F_max_ and p = 0.8306, r = 0.01 for duration), and a significant decrease in fluorescence onset times (Wilcoxon signed-rank tests, p = 5.72e-11, r = 0.30) over time (Fig. 3D–F). At the single-cell level, however, responses were heterogeneous: some neurons displayed progressively stronger responses to the same stimulation (Fig. 3G), while others remained relatively stable (Fig. 3H). By week 4, more neurons exhibited fluorescence intensities exceeding mean + STD (Fig. 3I), indicating enhanced excitability. To trace the source of this increase, we followed the subgroup of neurons with high ΔF/F_max_ in the final week back to the onset of training. These neurons showed significant increases in fluorescence over training, whereas the remaining neurons (blue) did not (two-sided Mann-Whitney U test; red: n_early_ = 52, n_late_ = 57, p = 9.76e-4, r = 0.32; blue: n_early_ = 174, n_late_ = 220, p = 0.1176, r = 0.08). The high intensity subgroup initially exhibited fluorescence comparable to the rest of the population (week 0: p = 0.5078, r = 0.0355; week 1: p = 0.0988, r = 0.0885) but diverged over time, showing significantly stronger responses by week 2 (p = 0.0018, r = 0.16) that persisted until the end of learning (week 3: p = 8.62e-15, r = 0.41; week 4: p = 2.48e-9, r = 0.31). This divergence coincided with the period during which behavioral detection thresholds decreased and stabilized (Fig. 3J).

To more accurately estimate spike timing and counts, we deconvolved Ca²⁺ transients to infer spike trains (Fig. 3K, Methods). As expected, the inferred spike trains mirrored the longitudinal changes in fluorescence intensity (Fig. 3L, M): neurons in the high intensity subgroup exhibited a significant increase in spike counts and a decrease in spike onset latency from early to late phases, while the remaining neurons exhibited little change. This subset initially showed spike counts and onset latency comparable to the rest of the population (Two-sided Mann-Whitney U tests; for spike counts week 0 - 1: p = 0.9699, r < 0.01; p = 0.1076, r = 0.08, respectively; and for onset latency week 0 – 1 : p = 0.8767, r < 0.01; p = 0.6763, r = 0.02, respectively), but diverged from week 2 to week 4 (for spike counts week 2 - 4: p = 0.0070, r = 0.14; p = 3.13e-14, r = 0.40; p = 1.16e-8, r = 0.30, respectively; and for onset latency week 2 – 4: p = 0.0442, r = 0.10; p = 0.0024, r = 0.16; p = 0.0059, r = 0.14, respectively). Collectively, these findings suggest that a subset of ICMS-activated neurons undergoes experience-dependent plasticity in response to repeated stimulation, reflected in increased excitability and reduced response latency over time, paralleling the time course of behavioral learning.

Lastly, we examined neurons newly recruited in the later weeks (Fig. 3N) — after detection thresholds had decreased and stabilized — and compared their responses to those of neurons consistently activated throughout training. Neurons recruited exclusively during the last week of training exhibited lower ΔF/F_max_ values (Kruskal-Wallis tests with Tukey-Kramer post-hoc correction, p **=** 3.09e-19; n_wk0_ = 23, n_wk2-4_ = 57, n_wk4_= 700) and fewer deconvolved spikes (p = 3.29e-17) than the high-intensity subgroup during weeks 2–4 while activation onset time remained similar between the two groups (p = 0.0633). Additionally, newly recruited neurons at week 4 exhibited lower fluorescence intensity (p = 0.0349) and shorter deconvolved spike onset (p = 0.479) than all tracked neurons at week 0 before learning-induced fluorescence enhancement occurred. Consistent with results in Fig. 3**D**-**H**, the enhanced subgroup at weeks 2-4 had significantly increased ΔF/F_max_, greater spike counts, and shorter spike onset time compared to all longitudinally tracked neurons at week 0 (ΔF/F_max_: p = 0.0114; spike count: p = 0.0059; spike onset: p = 0.0030). These results suggest that the late-recruited neurons were likely engaged indirectly, possibly through network-level mechanisms that emerged during learning.

### Chronic electrophysiology confirms learning-driven strengthening of neuromodulation

To capture fast neural dynamics beyond the temporal limits of two-photon Ca²⁺ imaging, we performed intracortical electrophysiological recordings from the same NET employed for ICMS, enabling direct measurement of spiking activity. Imaging and recording were conducted simultaneously and targeted complementary populations: two-photon imaging provided broad volumetric coverage, whereas electrophysiology sampled over 1 mm in depth across the full cortical column (Fig. 4A, B), recording a total of 973 units in 6 animals.

**Fig. 4.**
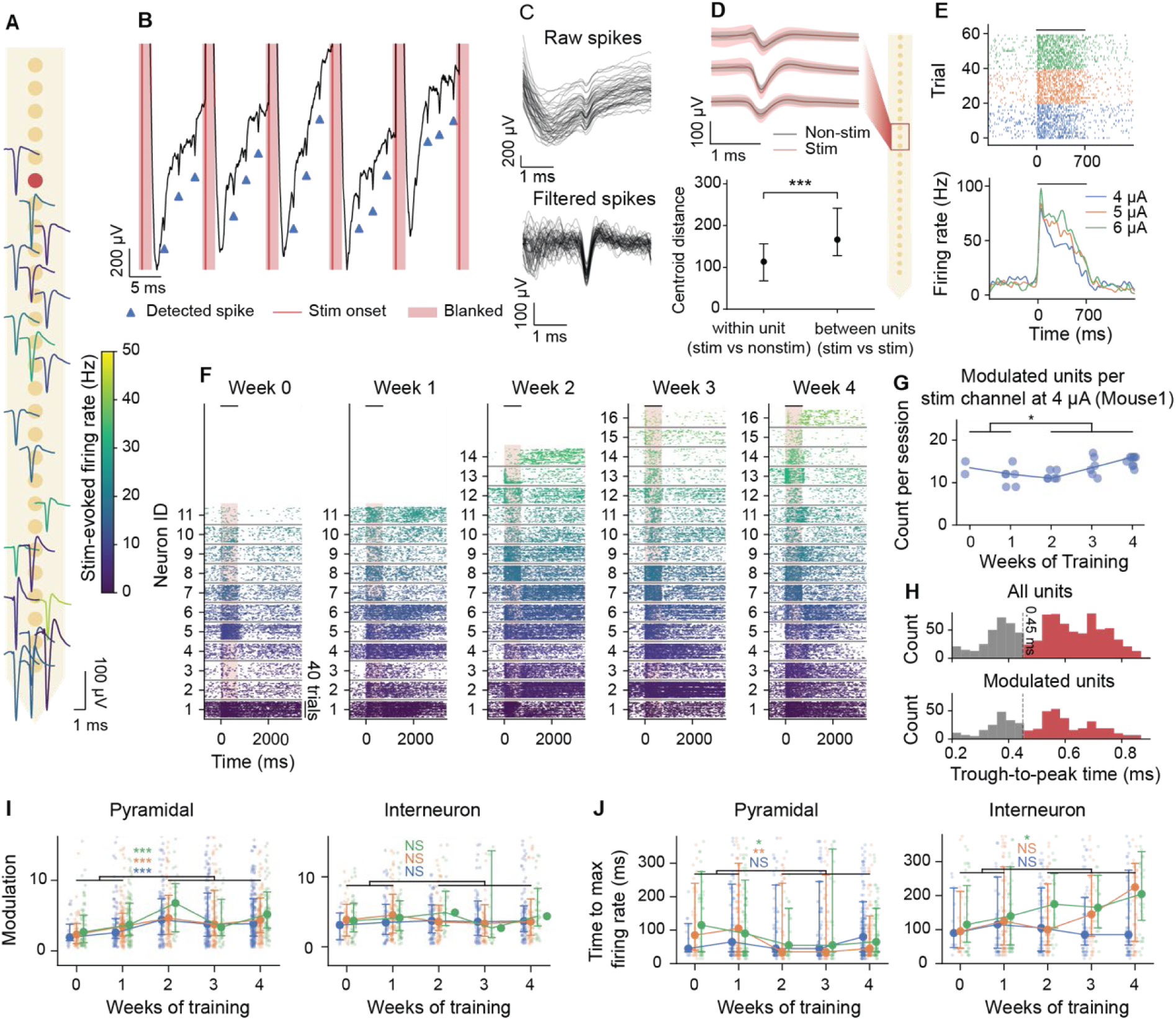
Intracortical recording reveals enhanced ICMS-evoked modulation over training. **(A)** Single-unit waveform overlaid on estimated recording locations. Red dot: stimulation contact. Color codes firing rate during ICMS. (**B**) Raw voltage trace and detected spikes (blue triangle) from a unit in (**A**) during ICMS at 100 Hz. Vertical lines mark stimulation and shading marks blanked out period (0.5 ms before and 1.5 ms window after each pulse). (**C**) Stimulation-obscured raw spike waveforms (top) and recovered waveforms (bottom) from a representative unit. (**D**) Top: example average waveforms of a neuron across three consecutive contacts during stimulation (red) and non-stimulation waveforms (black), showing waveform similarity. Dark line and shaded regions indicate mean ± STD. Bottom: Within-unit distances are significantly smaller than between-unit distances from nearby units in the PC space. (**E**) Spike raster (top) and trial-averaged firing rates (bottom) of a representative unit. Color indicates current amplitudes. (**F**) Spike rasters of all modulated cells in one animal over time (4 μA, one stimulation contact). Units are ordered by cortical depth and not tracked across sessions. In **E**, **F**, horizontal bars mark stimulation duration. (**G**) The number of modulated units of Mouse 1 at 4 μA over time. Dot represents a stimulation channel; line shows the median. (**H**) Bimodal distribution of waveform trough-to-peak intervals from all recorded units (top) and modulated units during stimulation (bottom). Dashed line at 0.45 ms marks the putative classification of narrow-waveform interneurons (gray) and pyramidal neurons (red). (**I, J**) ICMS-induced modulation (**I**) and time to maximum firing rate (**J**) of pyramidal cells and interneurons over weeks in training. Filled circles denote the median; error bars denote interquartile range (IQR). n = 6 animals for **H-J**. **D**, **I**, **J**: Two-sided Mann-Whitney U test; **G**: one-sided Mann-Whitney U test. NS, not significant, * p<0.05; ** p<0.01, and *** p<0.001.

Because stimulation artifacts can obscure evoked spikes and interfere with spike detection and sorting, we developed a custom spike-sorting pipeline with pre- and post-processing steps to suppress artifacts and recover waveforms for clustering (Methods, Fig. 4C). Principal-component (PC) analysis confirmed that stimulation-evoked waveforms closely matched non-stimulation waveforms and remained distinct from neighboring units (Fig. 4D, fig. S3A): within-unit Euclidean distances in the PC space (between stimulation and non-stimulation waveforms of the same unit; n = 243) are significantly smaller than the between-unit distances (between stimulation waveforms of different nearby units; n = 801; p = 1.6e-31, r = 0.49). This relationship was consistent across varying numbers of PCs (3, 5, 7, 10 [default], 15, and 20) indicating robustness to the choice of dimensionality.

Electrophysiology revealed rich temporal firing patterns of 484 neurons significantly modulated by the 700-ms stimulation train (paired t-test, p < 0.05, Methods), including strong onset activation followed by adaptation (Fig. 4E), features not resolvable with Ca²⁺ imaging. We quantified the modulation effect using a *modulation score*—the *z*-score of the average spike count during stimulation relative to the pre-stimulus baseline across trials and current amplitudes—and *activation latency*, defined as the time between stimulation onset to maximum firing rate. Representative longitudinal recordings (Fig. 4F) exhibited activation patterns consistent with Ca^2+^ imaging: average modulation scores of all cells increased over the course of training (fig. S3B), while activation latency decreased mildly (fig. S3C). When stimulation currents were adjusted to session-specific detection thresholds, the modulation score of these units remained stable across training (fig. S3D). Different from imaging results, which revealed a clear expansion of the activated population over time, electrophysiological recordings detected only a moderate increase in modulated unit numbers in some animals (Fig. 4F, G, n_early_ = 8, n_late_ = 21, p = 0.044, r = 0.42) and no significant group-level change, likely due to the sparser spatial sampling of electrophysiology.

While Ca²⁺ imaging in Thy1-GCaMP mice primarily captures excitatory pyramidal neuron activity(*39*), electrophysiology detects both excitatory and inhibitory neurons, which can be classified as putative pyramidal neurons or fast-spiking interneurons based on spike waveform and firing dynamics(*40*). Modulated units (n = 484; 311 pyramidal, 173 interneuron) showed a bimodal spike-width distribution similar to the full recorded population (n = 973; 672 pyramidal, 301 interneuron) (Fig. 4H), confirming robust spike detection on both cell types. Notably, increases in modulation strength and decreases in latency were driven primarily by pyramidal cells (modulation strength 4 μA: p = 2.6e-5, r = 0.25; 5 μA: p = 5.3e-5, r = 0.22; 6 µA: p = 2.0e-4, r = 0.23; latency 4 µA: p = 0.32, r = -0.06; 5 µA: p = 1.7e-3, r = -0.17; 6 µA: p = 0.049, r = -0.12), whereas interneurons showed little significant change (Fig. 4I–J, modulation strength 4 µA: n_early_ = 69, n_late_ = 254, p = 0.39, r = 0.07; 5 µA: n_early_ = 113, n_late_ = 180, p = 0.45, r = -0.05; 6 µA: n_early_ = 122, n_late_ = 86, p = 0.24, r = 0.10; latency 4 µA: p = 0.88, r = -0.01; 5 µA: p = 0.31, r = 0.07; 6 µA: p = 0.010, r = 0.21). Overall, at the coarse 700-ms time scale, electrophysiology paralleled imaging results in showing increased modulation with learning, while providing improved temporal precision and cell-type resolution.

### Millisecond pulse-locking dynamics differentiate learning-induced plasticity

Beyond capturing overall neural modulation during the 700-ms stimulation train, the millisecond resolution of electrophysiological recordings enables precise analysis of dynamics between individual pulses spaced merely 10 ms apart. To assess the temporal precision of evoked spikes, we aligned each neuron’s spikes to individual stimulation pulses (Fig. 5A, B), computed spike probability within each 10-ms interpulse interval (Fig. 5C), and quantified phase-locking using peak firing probability and temporal dispersion (Methods). ICMS-activated neurons segregated into two groups based on temporal alignment: pulse-locked (PL) responses, with spikes precisely aligned to each pulse (Fig. 5A–C, top), and non-pulse-locked responses, with spikes distributed throughout the interval (Fig. 5A–C, bottom). These distinct temporal patterns indicate different activation mechanisms: the precise alignment of spikes in PL neurons suggests direct or monosynaptic activation, while the broad distribution of NPL spikes reflects polysynaptic or network-mediated activation(*9*). We verified that spike detection worked well for both PL and NPL cells (fig. S4A).

**Fig. 5.**
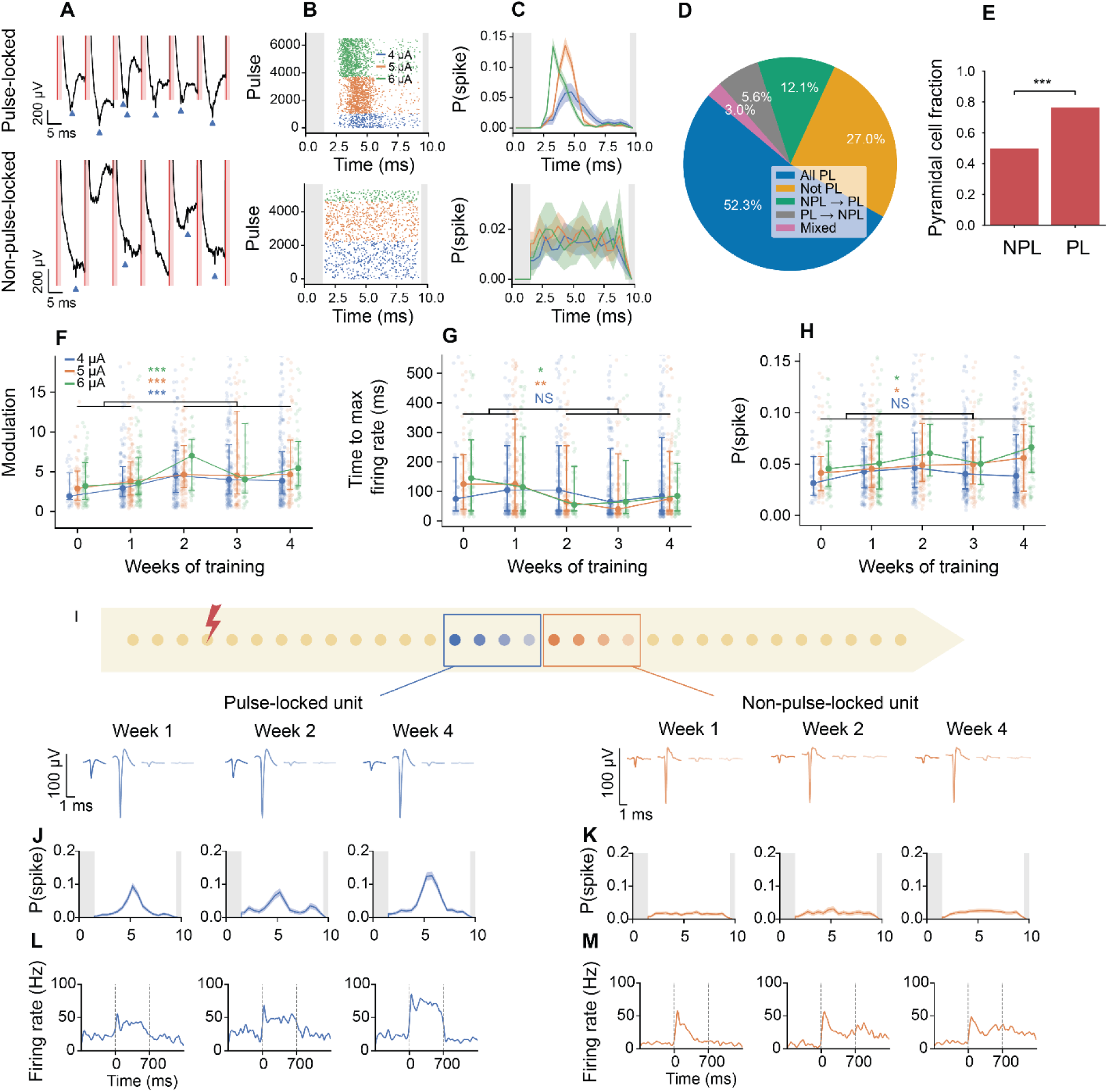
Distinct longitudinal changes of pulse-locked and non-pulse-locked responses during ICMS training. **(A**) Raw voltage traces and detected spikes (blue triangle) from representative pulse-locked (PL, top) and non-pulse-locked units (NPL, bottom) at 5 μA. Light red shading denotes blanking periods. (**B**, **C**) Spike raster (**B**) and spike probability (**C**) for the same units in (**A**) during the 10-ms interpulse intervals at 4, 5, and 6 μA. Solid lines and color-matched shading indicate means and STD. Gray shading denotes blanking periods. (**D**) Summary of pulse-locking stability across current amplitudes (4–6 µA) from the same stimulation site (*n* = 729 unit-channel pairs). Legend: “PL” – pulse-locked at all currents; “NPL” – non-pulse-locked at all currents; “NPL → PL” or “PL → NPL” – units switching classifications with increasing current; “mixed” – units showing both PL and NPL responses without current dependence. (**E**) Proportion of pyramidal cell type in PL units is higher than in NPL units. (**F, G**) Modulation (**F**) and onset time to max firing rate (**G**) of PL responses during 700 ms stimulation train at 4, 5, 6 µA from early to late phases. (**H**) Maximum spike probability within the interpulse interval of PL responses. (**I**) Representative tracking of two units over four weeks. Waveforms from a tracked PL and NPL unit recorded across weeks 1, 2, and 4 remained stable across both the primary and neighboring contact sites. Recording sites are overlaid on a NET probe diagram. (**J**, **K**) Maximum spike probability and firing rate (**L**, **M**) over time in training for a tracked PL unit (**J, L**) and a tracked NPL unit (**K, M**). n = 6 animals for **D**-**H**. Filled circles denote the median; error bars denote IQR. **E:** two-sided Fisher’s exact test; **F-H**: two-sided Mann-Whitney U test. NS, not significant, * p<0.05; ** p<0.01, and *** p<0.001.

Supporting these classifications, spike latency of PL responses increased linearly with distance from the stimulation site, consistent with action potential propagation along axons (fig. S4B). Linear fits to the monotonically increasing segment yielded conduction velocities of 0.2 – 0.27 m/s, in agreement with prior optical measurements in fluorescently labeled axons(*41*). Latency jitter, trial-to-trial variability in spike latency, did not vary with distance (fig. S4C), indicating stable spike timing of axonal conduction regardless of propagation distances. Because pulse-locking reflects the underlying activation mechanism, we expected these classifications to remain stable across stimulation currents. Indeed, over 80% of units retained their PL and NPL identity at a given stimulation contact as current amplitude increased (Fig. 5D). Pyramidal cells are more prevalent among PL units than NPL units (Fig. 5E, PL: 76.2%, NPL: 49.8%; two-sided Fisher’s exact test; p = 1.6e-9), consistent with the fact that narrow-waveform interneurons are often recruited indirectly through strong excitatory projections from local pyramidal neurons(*5*, *42–44*).

Importantly, PL and NPL responses exhibited distinct longitudinal changes during behavioral learning (n = 265 PL and 219 NPL units across 6 animals). At the coarse 700-ms train level, increases in modulation strength and decreases in latency were driven exclusively by PL responses (Fig. 5F, G), whereas NPL responses showed no significant changes (fig. S4D, E). Specifically, PL modulation increased from early to late phases across all current amplitudes (4 μA: n_early_ = 136, n_late_ = 469, p = 3.3e-5, r = 0.23; 5 µA: n_early_ = 188, n_late_ = 285, p = 2.4e-4, r = 0.20; 6 µA: n_early_ = 179, n_late_ = 135, p = 6.4e-5, r = 0.26), and latency decreased significantly at 5 and 6 μA (4 µA: p = 0.35, r = -0.05; 5 µA: p = 5.9e-3, r = -0.15; 6 µA: p = 0.040, r = -0.14). At the finer 10-ms interpulse scale, PL responses also showed increased peak spike probability over time at 5 and 6 μA (Fig. 5H, p = 0.22, r = 0.07; 5 µA: p = 0.028, r = 0.12; 6 µA: p = 0.010, r = 0.17). Because PL neurons are directly or monosynaptically activated, this enhanced modulation likely reflects increased axonal or somatic excitability, potentially through structural plasticity such as axonal enlargement(*45*) or strengthening of monosynaptic connections during learning.

Although we stimulated and recorded from S1, movement-induced widespread brain activity(*46*, *47*) could influence the neural response. To address the most direct concern regarding overt wheel movements, we used wheel movements as a proxy for actual movement, acknowledging that other movements (e.g., whisking, facial, or postural movements) could also occur. To limit movement confounds related to wheel movement, we restricted our analysis to an early post-stimulation window preceding movement onset. Wheel-movement onset, quantified on a trial-by-trial basis in all animals (n = 5) with reliable rotary encoder recordings (fig. S5A, B), decreased with training similarly to behavioral response times (fig. S1E, fig. S5C). We evaluated three candidate cutoffs from stimulation onset: the median wheel-movement onset in the fastest animal (Mouse 1, 122 ms; fig. S5D), the median onset across all hit trials (135 ms), and the median onset during the late training phase (129 ms), which was faster than in the early phase (fig. S5D). We selected the most stringent cutoff (the fastest animal median, 122 ms), rounded it down to 120 ms, and reanalyzed PL and NPL modulations within this restricted window across all 6 behavioral animals. This analysis yielded results consistent with those obtained using the longer 700 ms window (fig. S5E-H), supporting that the distinct longitudinal changes of PL and NPL cells arise from responses to ICMS rather than movement-related activity. Additionally, we extended a similar constrained time-window analysis to the two-photon imaging data, using the first five frames following ICMS onset (corresponding to approximately 132 – 165 ms). Any neurons whose fluorescence rose above the activation threshold after this time window were excluded (fig. S6A). Both recruitment of tracked neurons and the response amplitude of the increasing subset (Fig. 3J) increased significantly from early to late training (fig. S6B, C), consistent with the findings from the full response window. Furthermore, in the control group without behavioral learning, neither PL nor NPL cells showed significant changes in any of these metrics (fig. S7), further supporting that the longitudinal changes observed in the learning group are associated with behavioral learning.

To determine whether these distinct longitudinal differences arise from stable adaptations within individual neurons or shifts in the recorded population, we tracked single units over time using UnitMatch(*48*), a waveform matching algorithm that identifies the same neuron across recording sessions based on spatial position and waveform-based parameters (Methods). This yielded 34 tracked units spanning both early and late learning phases across 6 animals (fig. S8A, B). Fig. 5I shows two representative units, one pulse-locked and the other non-pulse-locked, tracked over the training period. Waveform distances between sessions of the same tracked unit were 6.8-fold smaller than distances to nearby (within 2 channels) non-matched units in the same session (p = 6.6e-83, Mann-Whitney U; fig. S8C), confirming reliable unit identification and tracking. The PL neuron exhibited increasing spiking probability within the 10 ms interpulse interval (Fig. 5J) and increasing modulation over the 700-ms pulse train (Fig. 5L), whereas the NPL neuron remained largely unchanged (Fig. 5K, M). Analyses of the 34 tracked units support the main findings: PL modulation increased significantly over training while NPL modulation did not, and PL time to maximum firing rate decreased (fig. S8D).

Together, the millisecond-resolved recordings revealed that behavioral learning engages two complementary forms of plasticity: strengthening of direct or monosynaptic excitation in PL neurons and expansion of network-level activity reflected in NPL neurons. Longitudinal tracking further demonstrates that these effects arise from stable, cell-specific adaptations, linking distinct activation mechanisms to the population-level plasticity underlying ICMS learning.

### Differential behavioral correlates of pulse-locked and non-pulse-locked neurons

To examine how activation mechanisms relate to behavior, we analyzed PL and NPL responses separately by behavioral outcomes (Fig. 6A). Representative units of each type exhibited distinct longitudinal changes: PL responses were similar between hit and miss trials, whereas NPL responses differed depending on outcome. Over time in training, the difference between hit and miss increased progressively for NPL units, suggesting that outcome-related differences strengthened with learning (Fig. 6A).

**Fig. 6:**
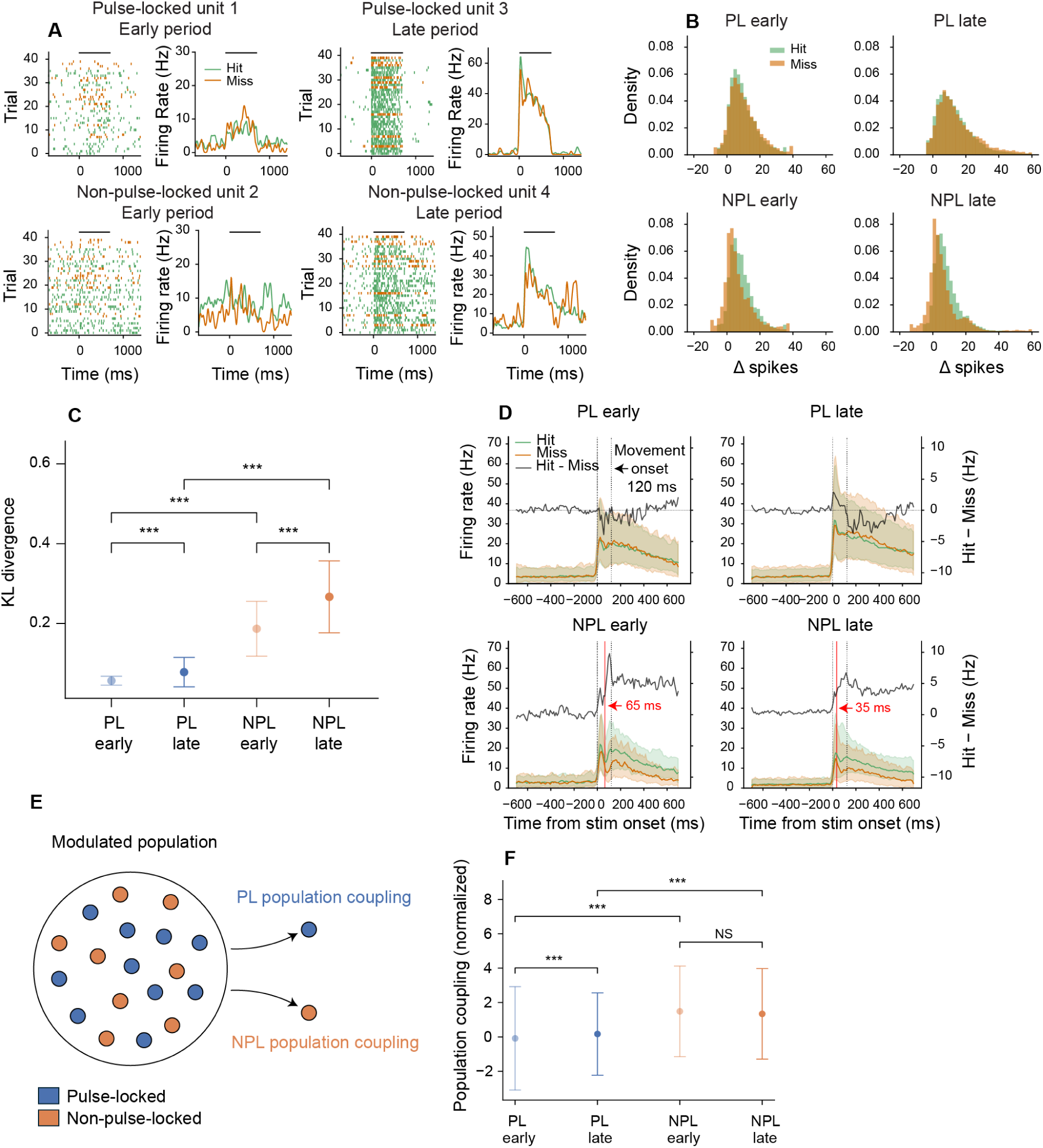
Pulse-locked and non-pulse-locked units exhibit distinct firing patterns linked to behavioral outcomes. (**A**) Example raster plots and trial-averaged firing rates of PL and NPL units from an early and late session (green: hit; orange: miss). All four units were recorded during 4 μA stimulation trials using the same stimulation channel in the same animal. (**B**) The distributions of Δspikes per trial across all animals and stimulation conditions in early and late phases. (**C**) Comparison of KL divergence of Δspikes distribution of PL and NPL responses at early and late phases. (**D**) Trial median responses of all PL and NPL units for hit (green) and miss (orange) trials. Shades represent IQR. The black lines plot the difference of median firing rates (Hit – Miss). Red vertical lines mark the onset time of significant divergence between hit and miss firing rate. (**E**) Schematic illustration of population coupling analysis, showing how stPR was calculated for PL (blue) and NPL (orange) units with respect to the modulated population. (**F**) Population coupling normalized to shuffled control of PL and NPL cells in early and late phases. Filled circles denote the median; error bars denote interquartile range. n = 6 animals. **C**, **F**: two-sided Mann-Whitney U with FDR-BH correction. NS, not significant, *** p<0.001.

To test whether this trend generalized across the population, we computed a Δspikes value for each trial, defined as the number of evoked spikes during the 700-ms stimulation train minus the number of pre-stimulation spikes in the preceding 700-ms window. Each trial inherited the pulse-locking classification of its corresponding unit (Fig. 6B). We quantified hit-miss differences in Δspikes distributions using Kullback-Leibler (KL) divergence, a measure from information theory(*49*), which quantifies how different two probability distributions are. Higher KL divergence indicates greater separation between hit and miss responses. While both PL and NPL separation between hit and miss trials increased over time (PL: p_FDR_ = 7.2e-26, r = 0.18; NPL: p_FDR_ = 6.1e-3, r = 0.05), NPL responses showed greater separation between hit and miss trials than PL responses in both early (p_FDR_ = 7.3e-116, r = 1.00) and late (p_FDR_ = 7.3e-116, r = 0.34) phases.

We next extended our analysis beyond the overall separation of the hit and miss responses to examine when they diverged significantly. We examined the trial-averaged firing rates of all units (median ± IQR) and computed the differential firing rate between hit and miss trials (Fig. 6D). Significant divergence was identified using cluster-based permutation testing(*50*), allowing us to determine both the presence and timing of these differences. Despite an increase in KL divergence from early to late learning, PL responses did not show any significant hit-miss divergence at any time point. In contrast, NPL responses diverged significantly beginning at 65 ms post-ICMS onset during the early period. With training, the onset of divergence in NPL neurons shifted earlier to 35 ms. Importantly, both onsets occur before movement onset (∼120 ms), suggesting that NPL neurons carry more behaviorally relevant information about sensory input, and that this encoding strengthens with training.

To further compare the integration of PL and NPL cells with local circuit, we computed population coupling, which measures how strongly a neuron’s activity co-varies with the surrounding population(*51*). To isolate coupling during stimulation, we calculated the spike-triggered population rate (stPR) between each modulated unit, PL or NPL, and the entire modulated population within the stimulation window (Fig. 6E). PL coupling increased over time (Fig. 6F, n_early_ = 494, n_late_ = 899, p_FDR_ = 1.95e-6, r = 0.16) while NPL coupling remained stable (n_early_ = 263, n_late_= 553, p_FDR_ = 0.33, r = -0.04). The increase of PL cells’ coupling to the entire population was driven primarily by stronger coupling among PL cells (fig. S9A), reflecting strengthened temporal coordination likely associated with their enhanced spike probability and reduction in firing latency. In both early and late learning phases, NPL units exhibited significantly higher population coupling than PL units (early: p_FDR_ = 3.4e-25, r = 0.46; late: p_FDR_ = 1.7e-22, r = 0.31), supporting their stronger network integration during stimulation and greater behavioral relevance. This difference remained robust when computed within the 120 ms pre-motion time window (fig. S9B) and was absent in control animals without behavioral learning (fig. S9C), supporting that it reflects learning-mediated responses to ICMS rather than nonspecific effects. Together, these findings reveal complementary roles of PL and NPL neurons in ICMS learning: PL cells undergo progressive adaptation of enhanced responsiveness to stimulation, whereas NPL cells exhibit network-level plasticity that more strongly links neural activity to behavioral outcomes.

## Discussion

Understanding the mechanisms of neural plasticity at single-neuron resolution during electrical-stimulation-induced learning has long been technologically prohibitive. Prior ICMS studies have largely focused on either behavioral outcomes such as detection thresholds and perceptual reports(*3*, *52–55*), or single-shot neural responses immediately after stimulation(*10–13*), without examining how neural activity reorganizes over chronic timescales as learning progresses. The key technologically enabling advance of this study is the integration of high-resolution optical and electrophysiological readout with stable ICMS and behavioral learning. This multimodal approach fills a critical knowledge gap by directly tracking how stimulation-evoked responses evolve over time as animals learn to associate an artificial sensory input with behavior.

Conceptually distinct from previous work, we find that ICMS-induced plasticity during learning is neither a uniform elevation of cellular excitability nor solely a Hebbian strengthening of networks(*56*). Instead, plasticity appears to depend on the activation mechanism and reflects a combination of both processes (Fig. 7). Neurons activated directly or monosynaptically exhibited changes consistent with strengthening of excitability, exhibiting increased spiking rate and reduced latency in response to stimulation. In contrast, neurons recruited indirectly showed expanded population recruitment, strengthened population coupling, and tighter correlation with behavioral performance, consistent with experience-dependent network changes. While brain-slice experiments(*57*, *58*) have shown rapid excitability changes in stimulated neurons during and immediately after stimulation, these preparations cannot capture how intact cortical circuits reorganize during behavioral learning. In vivo optogenetic studies have reported excitability changes over weeks when stimulation is paired with learning(*16*, *17*), but these studies relied on calcium imaging, which lacks the temporal resolution to distinguish direct and monosynaptic from network-mediated activation. Genetic modification may also alter intrinsic excitability(*58*), complicating interpretation. By complementing calcium imaging with electrophysiological recording of spiking activity, our results suggest that stimulation-induced plasticity may be organized according to activation mechanisms, providing a framework that unites excitability and connectivity-based processes into a single model of ensemble reorganization. The absence of comparable changes in the control group without behavioral learning further indicates that these effects are learning-dependent rather than driven by repeated stimulation alone, consistent with optogenetic studies showing that long-term plasticity requires behavioral engagement(*16*).

**Fig. 7.**
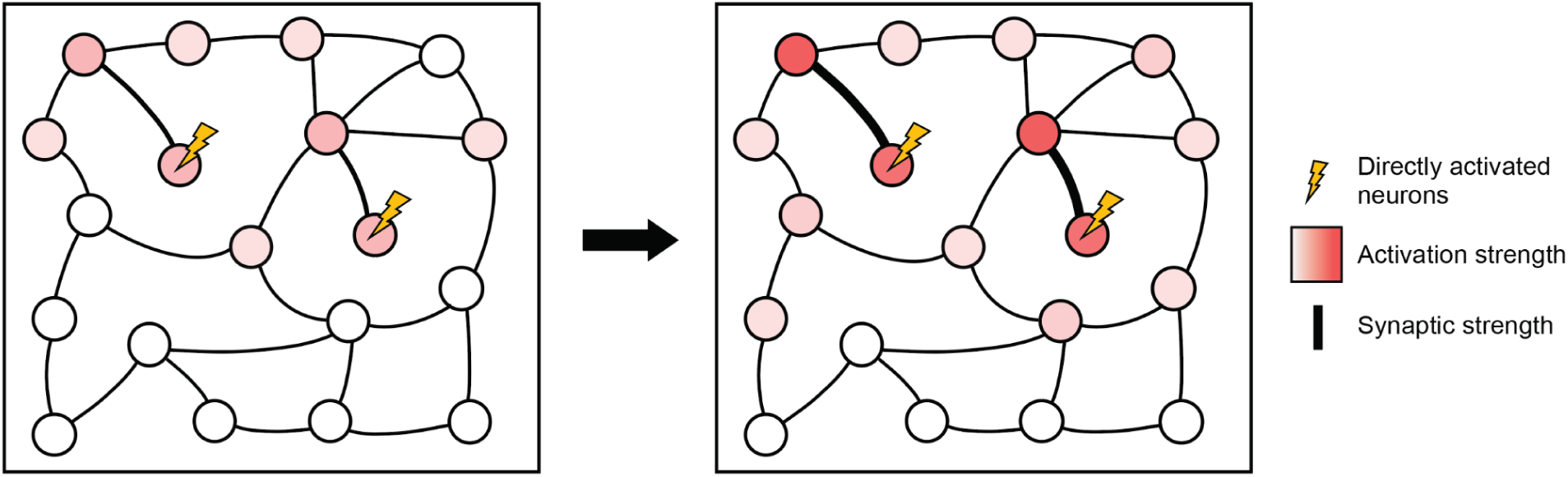
Schematic illustration of the proposed mechanism of the activation-mechanism-dependent plasticity. Neurons activated directly or monosynaptically increase their excitability (darker red shading) and strengthen monosynaptic connections (thicker connection lines) over training. In contrast, polysynaptically activated neurons maintain stable activation strength but show greater recruitment.

The enhancement of excitability in PL neurons may arise from two non-exclusive mechanisms. First, directly activated neurons may undergo axon initial segment (AIS) plasticity, altering the threshold for action potential initiation(*59*, *60*). Second, monosynaptic inputs to directly activated cells may strengthen over time, amplifying the pulse-locked responses. These mechanisms could contribute to the emergence of a more reliable artificial input, thereby facilitating broader recruitment of NPL neurons. Although NPL neurons did not exhibit changes in intrinsic excitability, their activity was more tightly linked to behavioral outcomes, consistent with plasticity in polysynaptic or recurrent network connections that may influence circuit output. Our findings that indirectly activated neurons are more behaviorally relevant and exhibit expanded population recruitment align with prior observations from optogenetic stimulation(*17*) and ICMS studies(*9*). Our results uniquely reveal divergent trajectories of directly and indirectly activated populations, suggesting a division of labor in which excitability and synaptic strengthening consolidate the fidelity of direct activation, whereas network-level plasticity links the artificial input to behavioral relevance. Because PL neurons are directly driven by stimulation, their responses are dominated by the input itself and therefore exhibit limited modulation by behavioral outcome. This is consistent with findings in optogenetic stimulation paradigms where directly-activated opsin-expressing neurons showed no differential activity between different behavioral trials, whereas downstream non-expressing neurons did(*17*).

Beyond distinguishing the mechanisms of plasticity in these two populations, our data also reveal additional features of neural activation underlying ICMS-induced learning. First, we observed a robust decrease in behavioral detection threshold over training when stimulation was delivered through ultraflexible electrodes that minimize confounds at the tissue-electrode interface. This finding helps reconcile previously inconsistent reports of longitudinal threshold changes with conventional electrodes, where thresholds have been shown to increase(*19*, *20*, *26*, *61*) and/or decrease(*3*, *21*, *23*)over time. By minimizing adverse tissue responses, such as neural degeneration, glial scarring, and encapsulation, which can elevate thresholds, our results suggest that learning contributes to a reduction in detection thresholds when interfacial stability is preserved. Second, sustaining the performance of a learned ICMS detection task required a consistent number of activated neurons at consistent modulation strength, even as detection thresholds shifted with training. This suggests that learning to process the artificial input does not reduce the neural resources required for perception but instead depends on a stable ensemble to maintain reliable detection. Third, we found a surprisingly large fraction of indirectly activated neurons when ICMS was paired with behavioral learning, much greater than in studies without a behavioral task(*5*). This difference may reflect the engagement of network recruitment during behavioral learning, an effect not captured by stimulation alone. Thus, studies of ICMS activation must consider how behavioral learning shapes neural recruitment.

Learning driven by ICMS shares key features with natural sensory learning but also exhibits important differences. In natural paradigms, such as whisker detection(*62*) and visual discrimination(*63*) tasks, learning typically enhances response reliability and selectivity within defined sensory representations, often without broad increases in population recruitment. In contrast, ICMS evokes spatially distributed activation patterns, and we observe an expansion of the recruited population at fixed currents alongside improved behavioral sensitivity during learning. Despite this difference, both forms of learning converge on the emergence of a stable, task-relevant neural ensemble, as reflected here by the consistent number of activated neurons at detection threshold despite decreasing thresholds over learning. The similarity and difference suggest that while artificial and natural learning may engage shared principles of experience-dependent plasticity, ICMS may rely more heavily on progressive network recruitment due to the non-specific nature of electrical activation.

Our findings have direct implications for the design of stimulation-based neuroprosthetics. A central challenge for sensory prostheses is not only to evoke reliable percepts but also to enable the brain to learn and connect behavioral meaning with artificial inputs. The reduction in detection thresholds with training indicates that experience improves sensitivity to artificial inputs when interfacial stability is maintained. At the same time, this plasticity implies that stimulation thresholds may need to be adjusted during learning, avoiding over-activation that could saturate responses and degrade resolution. Reliance on psychometric tests may be infeasible(*64*), especially for high-dimensional stimulation strategies(*65*). An adaptive stimulation paradigm, using real-time neural recordings to monitor recruitment and adjust stimulation levels accordingly, may provide a potential approach to maintain stability and flexibility, ensuring that artificial inputs remain both detectable and behaviorally meaningful as learning unfolds.

Several limitations should be considered when interpreting our results. First, the dissociation between PL and NPL neurons suggests distinct functional roles, but alternative interpretations remain possible, including that enhanced PL excitability drives downstream network recruitment reflected in NPL activity. While NPL responses are consistent with polysynaptic activation, indirect contributions from movement planning or state-related activity cannot be fully excluded despite our control analyses of a much shorter time window before wheel turning. Second, due to the limited imaging depth, we restricted stimulation to contacts located 200–400 μm below the cortical surface to ensure adequate sampling of neurons surrounding the stimulation site. As a result, we do not attempt to infer layer-specific differences in either neural responses or stimulation effects. Third, we use wheel movement as a proxy for behavioral output and do not fully capture body, head, or limb movements that may occur spontaneously or in response to stimulation. Because motor behaviors are accompanied by widespread cortical activity(*46*, *47*), and movement occurred within the 700 ms measurement window, learning-related changes in motor behavior could potentially contribute to the observed neural activity changes(66). Lastly, a substantial portion of the 700-ms analysis window occurred after the likely time of perceptual decision formation. Consequently, learning-related changes observed during this period may reflect not only stimulus-evoked sensory activity but also feedback and other behavioral-state-dependent signals.

## Materials and Methods

### Experimental model and subject participant details

A total of 12 mice were used in this study: 6 behavioral mice (4 male, 2 female) and 6 control mice (3 male, 3 female). All animals were C57BL/6J-Tg(Thy1-GCaMP6s)GP4.3Dkim/J, at least 8 weeks of age or older, and were bred on-site from breeding pairs acquired from Jackson Laboratories (Bar Harbor, ME). All 12 mice underwent NET implantation; however, the 6 control mice, which were part of a previous study(*22*), did not participate in behavioral training. Control animals underwent similar stimulation conditions, including stimulation parameters and longitudinal experimental timelines, and were subject to simultaneous imaging and recording as in the behavioral group. No behavioral apparatus such as a wheel or licking port was used for the control group. Mice were single-housed following implantation of NETs in the Animal Resource facility at Rice University. All surgical and experimental procedures in this study were in compliance with the National Institutes of Health Guidelines for the Care and Use of Laboratory Animals and were approved by the Rice University Institutional Animal Care and Use Committee (protocol IACUC-25-188).

### NET fabrication

In this study 32-channel, single-thread stimulating NETs were fabricated and implanted in animals for longitudinal behavioral testing. The NET fabrication has been previously described(*22*) but is briefly described here for clarity. Conventional photolithography and metallization on fused-silica wafers are used to fabricate the multi-layer structure of the NET electrodes. The microfabrication procedure was performed in the following steps. i) A nickel metal release layer was patterned by depositing 3 nm Ti and 60 nm Ni under the flexible section of the device. ii) An insulation layer was deposited on both the top and bottom surfaces using spin-coated, diluted polyimide (PI2574, HD Microchemicals) to a thickness of 500 nm. The coating and curing process was performed twice, resulting in a total insulation thickness of 1 µm. iii) An interconnect layer was defined by photolithography and metallization of a 3 nm Cr, 100 nm Au, and 3 nm Cr metal stack by electron (e)-beam evaporation (Sharon Vacuum Co., Brockton, MA). Layers of 3 nm Cr, 160 nm Ni, and 80 nm Au were deposited on the solder pads to improve solder reflow reliability and reduce alloying between solder and gold. iv) The top insulating layer was deposited to a thickness of 500 nm using the same method as the bottom layer. v) The thread outline, via to the electrodes, and solder pads were defined by RIE etching (Oxford Instrument) using an O2/CF4 gas mixture in the 9:1 ratio. vi) Microcontacts for recording and stimulation were defined by photolithography and sputter coating of 10 nm Ti, 100 nm Pt, 10 nm Ti, and 300 nm IrOx stack (AJA ATC Orion Sputter System). vii) Low-temperature solder balls were placed on solder pads to form a ball grid array using a solder jetting tool (PacTech), and the wafer was diced into individual devices. viii) The NETs were then individually bonded to custom printed circuit boards to interface with recording/stimulation electronics. ix) The flexible section of NETs was released from the substrate by etching the Ni layer, and the glass substrate was cleaved to the desired length. x) Lastly, the flexible implantable portion of the NET was hooked onto a 50-µm diameter sharpened tungsten wire and adhered via Polyethylene glycol (PEG), which served as a temporary adhesive securing the probe for implantation.

### Electrical stimulation and ephys recording system

Electrical stimulation was delivered through the Pico32+Stim front end and Grapevine Neural Interface Processor (Ripple Neuro, Salt Lake City, UT) to NET contact sites with impedances less than 1 megaohm at 1 kHz. Stimulation consisted of cathode-leading biphasic pulses with a 167 μs phase duration and 67 μs interphase interval, delivered at 100 Hz. Both stimulation and recording were performed via the Pico32+Stim front end, which was controlled using the Xippmex MATLAB API (MathWorks) on a computer separate from the behavioral task controlling computer.

### Surgical procedure

All animals received co-implantation of a cranial optical window and NET in one surgery. Briefly, animals were anesthetized with isoflurane (3% for induction and 1%–2% for maintenance) and administered extended-release Buprenorphine (Ethiqa TM) and Dexamethasone (2 mg/kg, SC) for analgesia and to reduce surgery-induced inflammation, respectively. The surgical site was infiltrated with lidocaine (7 mg/kg 0.05%) subcutaneously before shaving and disinfected with iodine and alcohol wash before the initial incision into the skin of the head. The skull was exposed between the bregma and lambda skull sutures, followed by removing the fascia and scoring the skull crosshatch pattern to prepare the skull. A circular craniotomy 3 mm in diameter over the somatosensory cortex was drilled in the skull for the StimNET implantation, and a burr hole was drilled in the contralateral hemisphere to accommodate a Type 316 stainless steel grounding wire. Following the opening of the craniotomy, a 32-contact NET, affixed to a 75 µm tungsten wire using a bio-dissolvable PEG adhesive, was implanted through the dura into the somatosensory cortex using stereotaxic targeting approximately 2 mm ML and −1.5 mm AP at an insertion angle of 30° from vertical. Minor adjustments to the target coordinates were made to avoid surface vasculature and to ensure a clear region around the implantation site for imaging. As a result, the final distribution of implantation locations was ML: 1.787 ± 0.507 mm and AP: −1.533 ± 0.295 mm across 12 animals. Following the implantation of the NET, the PEG affixing the NET to the shuttle wire was allowed to dissolve, and the wire was removed. A sterile 4 mm glass coverslip window was secured over the craniotomy using cyanoacrylate adhesive and Metabond dental cement (Parkell, NY), with regions not directly covered by glass filled with Kwiksil (World Precision Instruments). Additional dental cement was applied to adhere a headbar for head fixation to the skull cap and seal the cranial window to the skull. Animals were provided with at least three days of recovery post-surgery and an additional three days of familiarization with the head restraint before the beginning of any experiments. The depth of implantation was targeted via stereotaxic linear drive. The insertion was prompt to prevent premature detachment between PEG and NETs. Rice IACUC approved all the animal experiments.

### Behavioral task

Mice were trained to perform an intracortical microstimulation (ICMS) detection task requiring them to turn a wheel past an angular threshold to receive a 3 μL water reward (Fig. 1B). Each trial consisted of four periods: a randomized pre-stimulus period (1-1.5 s), a stimulus-response window (700 ms), a fixed post-stimulation period (2.5 s), and a randomized intertrial interval (1-1.5 s). During the pre-stimulus period (PSP), mice were required to withhold movement such that any premature turns reset the PSP timer, delaying stimulus onset. In the stimulus period, mice received 100 Hz stimulation for the full 700 ms duration (70 pulses total). Mice were trained to make a “Go” response by turning the wheel past an angular threshold during the stimulation window, which immediately triggered a reward without terminating the stimulation. No stimulation was applied during the post-stimulation period and intertrial interval. To minimize photobleaching during calcium imaging, imaging was restricted to the time between the start of the PSP and the end of the stimulus-response window (Fig. 1B). If the mouse remained in the PSP state for longer than 5 s, imaging was stopped and the trial transitioned to the intertrial interval.

The behavioral task was performed on a standardized experimental rig developed by the International Brain Laboratory(*67*) with modifications based on our previous study(*22*). Each mouse had three stimulation contact sites selected during behavioral shaping, which were used consistently across sessions. Stimulation amplitudes were varied using a method of constant stimuli. Before each session, a brief amplitude sweep was performed to estimate the 50% detection threshold for each site. Sessions were structured into four blocks of 100 trials each. Each block contained ten randomly interleaved conditions: 10 catch trials (0 μA) and 90 stimulation trials (3 channels × 3 amplitudes × 10 trials). Behavioral performance was monitored in real-time, allowing stimulation amplitudes to be adjusted between blocks to maintain performance near threshold.

The behavioral task was controlled by a BPod state machine (Sanworks) running on a dedicated computer. Transitions between behavioral states in the state machine were communicated to the Ripple Neural Interface Processor via TTL pulses through the digital I/O front-end, allowing synchronization of stimulation onset and event time logging on the main recording computer.

The behavioral shaping is similar to the method used previously(*22*). Briefly, after a post-surgical recovery period of 4 weeks to ensure tissue-probe interface stability(*36*), animals were placed on water restriction while undergoing habituation to the behavioral rig. After habituation, mice began the shaping protocol, during which they learned to turn the wheel to obtain a water reward. Mice were then required to turn the wheel in response to suprathreshold ICMS for reward. After animals produced consistent low-latency responses within 700 ms with a low PSP violation rate (<10% of trials) at suprathreshold ICMS, they graduated to detection threshold measurements.

During detection threshold measurements, sets of three currents were presented to the animals, and probability response curves were fitted to their responses to determine their 50% detection threshold. This procedure was run for each of the selected stimulation sites in a randomized manner. During each measurement session, four blocks of behavioral trials were presented to animals, each consisting of 100 trials during electrophysiological recording and volumetric two-photon Ca^2+^ imaging. For the first block of trials in a session, animals were presented with stimulation currents at their detection threshold from their prior session, as well as 1 μA above and below their prior threshold. Following the block of trials, if the animal detection threshold was not closest to the middle of the three currents provided, the three currents tested were adjusted in the following block to provide stimulation at the detection threshold and 1 μA above and below. Electrophysiological recordings were continuously collected during each block, and imaging was performed 1-1.5 s prior to stimulation onset until 2.5 s after stimulation offset. A fifth block was presented for planar imaging; this block was not used for electrophysiological analysis. Rice IACUC approved all the animal experiments.

### Analysis of behavioral data

Detection thresholds were estimated by fitting a cumulative Gaussian psychometric function to the proportion of hits at each stimulation amplitude for each channel, using the *python-psignifit 4.3* package(*68*, *69*). Thresholds were calculated independently for each block and defined as the stimulus amplitude corresponding to 50% detection. The false-positive rate for each block was calculated as the proportion of responses during the ten catch trials and was applied to the psychometric curves for all three stimulation channels. A session-level threshold for each channel was computed as the average of the four block-level thresholds. Only data that included 4, 5, and 6 μA stimulation amplitudes were analyzed since these were the most consistently presented across animals and sessions.

### Spike sorting during stimulation

Ephys data were captured at a 30 kHz sampling rate. To mitigate ICMS stimulation artifacts and imaging scanning artifacts, a custom preprocessing pipeline implemented in Python using the *SpikeInterface* package was applied to the raw data. First, each stimulation pulse was blanked or replaced with zeros from pulse onset to 1.4 ms after onset. We assume that the artifacts from the stimulation pulses are similar, given the same stimulation condition (stimulation channel and current). Therefore, we subtract the median of all post-stim segments across all pulses in a trial, where a trial constitutes 70 pulses of the same stimulation channel and current, from all pulse windows. Third-order polynomial fits were used to detrend each channel trace. Common referencing was applied. Cubic interpolation between 0.5 ms before onset and 1.4 ms after stim onset and a 300 Hz to 5000 Hz bandpass filter was applied. Traces were blanked from 0.5 ms before onset and 1.5 ms after onset.

The preprocessed data were then spike sorted using Mountainsort5 within the *SpikeInterface* framework. Following spike sorting, both automated and manual cluster curation were performed. Despite the preprocessing, residual contamination was observed in accepted clusters, necessitating a waveform curation step using a post hoc classification method (see code repository for details). To evaluate sorting accuracy, we assessed waveform separation during stimulation and non-stimulation periods (Fig. 4D, bottom). Each waveform was represented as a vector of size (number of functional channels) × 90 samples (3 ms). The waveforms were projected onto the first 10 principal components, and in this reduced PCA space, we computed the Euclidean distance between the centroids of unit i’s stimulation and non-stimulation spikes (within-unit distance) and between the stimulation centroids of units i and j (between-unit distance). Comparisons were restricted to neighboring units with estimated locations within 240 µm or recorded on electrodes separated by no more than four contacts.

### Quantifying neuromodulation metrics via electrophysiology

Processed electrophysiological data were organized into responses, defined as the activity of a single unit in a specific stimulation channel-amplitude condition, so that a particular unit could contribute to multiple responses. A response was classified as modulated if a paired t-test comparing spike counts during stimulation and pre-stimulus for all stimulation condition trials yielded a significant increase (p < 0.05), included at least 50 spikes, and an average baseline firing rate of at least 0.5 Hz.

The modulation score of a response was calculated as the z-score, computed by subtracting the mean pre-stimulation period firing rate from the mean firing rate during stimulation and dividing by the standard deviation of pre-stimulation firing rates across trials. To determine the probability of evoking a spike from a single stimulation pulse, spikes in the 10-ms interpulse window were binned into 0.5-ms bins. Blanking was applied to the first 1.5 ms and last 0.5 ms of each interpulse interval. Probability of spiking was therefore the number of spikes per bin normalized by the number of trials for a specific stimulation condition. A unit was defined as pulse-locked to a stimulation condition using criteria described previously(*9*). In brief, if the spike probability curve during the 10-ms interpulse interval had a pulse-locked index (PLI) above 99% of shuffled data, the null distribution of PLIs, the unit was labeled as pulse-locked. Mean and standard deviation of the spike-probability curves (Fig. 5C) were estimated using 5000 bootstrap resamples, each resampling 20% of trials with replacement.

Pulse-locked and non-pulse-locked responses were subsets of the modulated responses. For the analysis in Fig. 5E, a unit was classified as pulse-locked using a two-step procedure. First, for each stimulation current, the unit was labeled as pulse-locked at that current if it was classified as pulse-locked at the majority of stimulation channels. Second, the unit was classified as pulse-locked overall if the majority of currents were pulse-locked.

### Unit tracking via electrophysiological recording

Longitudinal unit tracking for electrophysiological recording was performed using UnitMatchPy(*48*) which matches neurons across recordings based on spatial position and waveform-based parameters. For each animal, average waveforms were extracted from SpikeInterface sorting analyzers for all curated units and formatted as concatenated multi-channel templates. UnitMatch computes six waveform similarity metrics (amplitude, spatial decay, waveform shape, centroid position, centroid volatility, and centroid trajectory) and applies a naive Bayes classifier to estimate the probability that units from different sessions represent the same neuron. Units with cross-session match probability >0.75 were linked into tracks using connected-component analysis. When multiple units from the same session were assigned to the same identity, the best-matching unit was retained. Tracks were refined by requiring a consecutive-session match probability ≥ 0.5 and only tracks spanning ≥ 3 sessions were included. Tracks with median pairwise waveform distance > 3× the global median were excluded, as were individual sessions with waveform distance > 5× the track median. This yielded 34 tracks spanning both early and late learning phases across 6 animals. Tracking quality was validated by comparing Euclidean distances of concatenated multi-channel waveforms(*29*) between tracked pairs and nearby non-matched units within two electrode contacts, yielding a 6.8-fold separation ratio (fig S8C).

### Population coupling

Population coupling for each unit was calculated as described previously with minor modification in the normalization procedure(*51*). Population coupling was computed for pulse-locked (PL) and non-pulse-locked (NPL) units during ICMS-evoked periods. For each stimulation condition (current × channel), spike trains were segmented to include only spikes occurring within stimulation trains, concatenated across trials, and converted to continuous time. A population firing-rate trace was constructed by binning (1 ms), Gaussian-smoothing (σ = 10 ms), and mean-centering the activity of all modulated units except the unit of interest. For each neuron, the spike-triggered population rate (stPR) was obtained by averaging the population trace within a ±100 ms window around each of its spikes, and population coupling (PC) was defined as the stPR value at zero lag. To enable cross-session comparisons, PC values were normalized to a circular-shuffle null distribution in which the population-rate trace was randomly phase-shifted relative to the neuron’s spike train while preserving its temporal statistics; normalized PC was computed as the observed PC divided by the median of 200 absolute shuffled PC values. Units with fewer than 50 spikes in stimulation periods were excluded from analysis. Population coupling was additionally computed using only spikes within the first 120 ms of stimulation for the movement control analysis of behaviorally trained animals (fig. S9B) and using the full stimulation window for stimulation-only control animals (fig. S9C).

### Two-photon imaging

Two-photon (2P) imaging was performed using a laser scanning microscope (Ultima 2Pplus Bruker, MA) equipped with a 16x water immersion objective (numerical aperture of 0.8, Nikon, NY) and an ultrafast laser tuned to 920 nm for fluorescence Ca^2+^ excitations (InSight X3, Spectra-Physics). Images with a resolution of 512 x 512 pixels of a field of view up to 1 mm x 1 mm were acquired at 30 fps using galvo-resonant scanners. After initial habituation and behavioral shaping, z-stack 2P imaging (25 µm per scan, 300 µm in depth span) was performed during each behavioral session over the course of training, during which head-restrained mice performed the behavioral task described above. During each trial of the behavioral tasks, volumetric or planar imaging was initiated at PSP onset which is 1-1.5 s prior to the onset of stimulation, 700 ms during stimulation, and continued for 2.5 s post the cessation of stimulation providing quantification of the neural activity, before, during, and after stimulation. Stimulation and 2P imaging were synchronized via a custom MATLAB script and TTL signals generated by PulsePal (Sanworks, NY) unit. Volumetric imaging was continually looped after a full image stack was acquired for the duration of each trial. For planar imaging, the imaging plane was fixed at 100 μm below the cortical surface, and frames were continually collected at 30 fps for the trial’s imaging duration.

### Detection and quantification of ICMS-evoked Ca²⁺ activation

ICMS-evoked neuronal activation was identified from both volumetric and single-plane two-photon Ca²⁺ imaging using a unified segmentation and threshold-based detection workflow. For volumetric imaging, raw stacks were aligned to trial structure using behavioral timestamps and averaged into pre- and mid-stimulation 3-D volumes. Baseline fluorescence statistics were estimated by pooling all pre-stimulation volumes from valid trials to generate voxel mean and standard-deviation (STD) maps. Session-level ΔF and variance maps were then computed to identify regions consistently modulated by stimulation.

Candidate neurons were segmented by applying 3-D iterative thresholding (FIJI/ImageJ) to the baseline STD, variance, and ΔF maps, followed by mask merging and 3-D connected-component extraction in MATLAB. ROIs smaller than 5 µm radius were excluded. For each ROI, ΔF volumes for mid-stimulation epochs were obtained by subtracting the baseline pre-stimulation volume. Voxels exceeding two times of their baseline STD were classified as significantly responsive. An ROI was considered activated on a given trial if ≥50% of its voxels crossed threshold during the stimulation window. Activated neuron counts for a given channel and current amplitude were computed as the number of ROIs meeting this criterion. Spatial activation profiles were quantified by measuring each activated ROI’s 3-D Euclidean distance from the stimulation site. The enclosing radius was defined as the median distance of activated neurons for that condition. Activation density was computed by binning ROIs into concentric spherical shells (100-µm increments) and dividing the number of activated neurons by shell volume. For visualization, densities were normalized across animals and weeks by the 99th percentile of all shell values. For single-plane imaging sessions, neuronal ROIs were extracted using EXTRACT(*70*, *71*) with a lowered SNR threshold (≥1) to capture low-amplitude ICMS responses. ROIs were post-processed to remove small or duplicated components, and per-trial fluorescence traces were computed analogously to volumetric ROIs. For single-plane data, an ROI’s response was classified as activated if its trial-averaged fluorescence within the stimulation window exceeded the mean + 2 SD of that ROI’s full trace (all pre-, mid-, and post-stimulation segments concatenated across valid trials), for a continuous run of >5 frames (∼165 ms) with onset before 700 ms.

### Deconvolution

Raw calcium traces from planar imaging data were deconvolved using the OASIS algorithm(*72*) with an AR(1) calcium decay model. Model parameters and the minimum spike size threshold were optimized per ROI, and small events were thresholded to obtain a sparse estimate of the underlying spike train. Deconvolved spike trains were then passed to a custom activation-extraction pipeline which aligned ΔF/F and spike traces to ICMS onset, averaged across valid trials, identified activated neurons using ROI-specific normalized thresholds, and computed activation metrics including onset latency, peak time, activation duration, volume above threshold, and deconvolved spike onset and count.

### Alignment and unit tracking via Ca^2+^ imaging

Longitudinal cell tracking was performed across weekly single-plane imaging sessions. Each session’s average pre-stimulus image was aligned to a reference session using PatchWarp(*38*), a block-wise nonrigid image registration framework. Registration was performed using a global Euclidean transformation followed by a local affine warp with a block size of 6 and 25% block overlap. For each session, the alignment was run 3 times and the transformation with the lowest mean absolute pixel error relative to the reference was retained. The resulting deformation fields were used to transform each session’s ROI masks into the reference coordinate frame. ROIs were matched across sessions by maximizing pixel-wise mask overlap and sets of ROIs appearing in ≥N sessions (N = 4 for Mouse1; N = 3 for Mouse2 and Mouse3) were defined as longitudinally tracked units. Longitudinal activation rasters and ΔF/F-derived response metrics were then computed for all tracked neurons. Alignment quality was evaluated by quantifying cross-session spatial intensity correlations across animals (n = 16 sessions across 3 animals), yielding a correlation (r = 0.904 ± 0.068, mean ± std), indicating reliable alignment across sessions.

### Statistics

Statistical analyses for imaging data (Fig. 2-3, fig. S2) were performed in MATLAB (MathWorks, MA). Statistical analyses for ephys data (Fig. 4-6, fig. S3-9) were performed in Python. Results and details of the statistical comparisons performed in the study were reported in the results section and figure legends. In this study, p < 0.05 was accepted as statistically significant. Summary statistics are reported as median ± interquartile range (IQR) because data distributions were non-normal (Shapiro–Wilk test, p < 0.05). The Mann-Whitney U-test was performed to compare metrics between early and late phases with either two-sided or one-sided tests. Multiple comparisons were corrected for by using Benjamini and Hochberg FDR correction. Fisher’s exact test was used with two-sided tests. Comparisons of neuronal subgroups (Fig. 3N) were performed using Kruskal-Wallis tests with Tukey-Kramer’s post-hoc correction.

## Acknowledgements

We thank the Animal Resource Facility at Rice University for animal care and housing, and the Rice nanofabrication facility for support on the microfabrication of StimNET.

## Author contributions

Conceptualization: LL, RK, RL

Methodology: LL, CX, RK, RL

Investigation: RK, RL, PZ, JM, CX, LL

Visualization: RK, RL, LL

Supervision: LL, CX

Writing — original draft: RK, RL, LL with input from CX

Writing — review & editing: RK, LL with input from RL

## Data and materials availability

Processed imaging and electrophysiological datasets supporting this study are available at https://doi.org/10.5281/zenodo.17727762. Custom analysis code used in this study is archived on Zenodo (Version v1.0, DOI: 10.5281/zenodo.19561006). The source repository is additionally available at: https://github.com/XieLuanLab/icms-activation-plasticity. This study did not generate new materials.

## References

1. S. J. Bensmaia, L. E. Miller, Restoring sensorimotor function through intracortical interfaces: progress and looming challenges. Nat. Rev. Neurosci. 15, 313–325 (2014).

2. M. E. Urdaneta, N. G. Kunigk, F. Delgado, S. I. Fried, K. J. Otto, Layer-specific parameters of intracortical microstimulation of the somatosensory cortex. J. Neural Eng. 18, 055007 (2021).

3. C. L. Hughes, S. N. Flesher, J. M. Weiss, J. E. Downey, M. Boninger, J. L. Collinger, R. A. Gaunt, Neural stimulation and recording performance in human sensorimotor cortex over 1500 days. J. Neural Eng. 18, 045012 (2021).

4. M. C. Dadarlat, R. A. Canfield, A. L. Orsborn, Neural Plasticity in Sensorimotor Brain–Machine Interfaces. Annu. Rev. Biomed. Eng. 25, 51–76 (2023).

5. M. H. Histed, V. Bonin, R. C. Reid, Direct Activation of Sparse, Distributed Populations of Cortical Neurons by Electrical Microstimulation. Neuron 63, 508–522 (2009).

6. S. Butovas, C. Schwarz, Spatiotemporal Effects of Microstimulation in Rat Neocortex: A Parametric Study Using Multielectrode Recordings. J. Neurophysiol. 90, 3024–3039 (2003).

7. A. S. Tolias, F. Sultan, M. Augath, A. Oeltermann, E. J. Tehovnik, P. H. Schiller, N. K. Logothetis, Mapping Cortical Activity Elicited with Electrical Microstimulation Using fMRI in the Macaque. Neuron 48, 901–911 (2005).

8. L. E. Osborn, D. P. McMullen, B. P. Christie, P. Kudela, T. M. Thomas, M. C. Thompson, R. W. Nickl, M. Anaya, S. Srihari, N. E. Crone, B. A. Wester, P. A. Celnik, G. L. Cantarero, F. V. Tenore, M. S. Fifer, “Intracortical microstimulation of somatosensory cortex generates evoked responses in motor cortex” in 2021 10th International IEEE/EMBS Conference on Neural Engineering (NER) (IEEE, Italy, 2021; https://ieeexplore.ieee.org/document/9441123/), pp. 53–56.

9. N. D. Shelchkova, J. E. Downey, C. M. Greenspon, E. V. Okorokova, A. R. Sobinov, C. Verbaarschot, Q. He, C. Sponheim, A. F. Tortolani, D. D. Moore, M. T. Kaufman, R. C. Lee, D. Satzer, J. Gonzalez-Martinez, P. C. Warnke, L. E. Miller, M. L. Boninger, R. A. Gaunt, J. L. Collinger, N. G. Hatsopoulos, S. J. Bensmaia, Microstimulation of human somatosensory cortex evokes task-dependent, spatially patterned responses in motor cortex. Nat. Commun. 14, 7270 (2023).

10. J. T. Sombeck, J. Heye, K. Kumaravelu, S. M. Goetz, A. V. Peterchev, W. M. Grill, S. Bensmaia, L. E. Miller, Characterizing the short-latency evoked response to intracortical microstimulation across a multi-electrode array. J. Neural Eng. 19, 026044 (2022).

11. Y. Hao, A. Riehle, T. G. Brochier, Mapping Horizontal Spread of Activity in Monkey Motor Cortex Using Single Pulse Microstimulation. Front. Neural Circuits 10 (2016).

12. R. Yun, J. H. Mishler, S. I. Perlmutter, R. P. N. Rao, E. E. Fetz, Responses of Cortical Neurons to Intracortical Microstimulation in Awake Primates. eneuro 10, ENEURO.0336-22.2023 (2023).

13. D. Youssef, J. H. Wittig, S. Jackson, S. K. Inati, K. A. Zaghloul, Neuronal Spiking Responses to Direct Electrical Microstimulation in the Human Cortex. J. Neurosci. 43, 4448–4460 (2023).

14. K. Runge, C. Cardoso, A. De Chevigny, Dendritic Spine Plasticity: Function and Mechanisms. Front. Synaptic Neurosci. 12 (2020).

15. L. Carrillo-Reid, W. Yang, Y. Bando, D. S. Peterka, R. Yuste, Imprinting and recalling cortical ensembles. Science 353, 691–694 (2016).

16. B. Akitake, H. M. Douglas, P. K. LaFosse, M. Beiran, C. E. Deveau, J. O’Rawe, A. J. Li, L. N. Ryan, S. P. Duffy, Z. Zhou, Y. Deng, K. Rajan, M. H. Histed, Amplified cortical neural responses as animals learn to use novel activity patterns. Curr. Biol. 33, 2163–2174.e4 (2023).

17. R. Pancholi, L. Ryan, S. Peron, Learning in a sensory cortical microstimulation task is associated with elevated representational stability. Nat. Commun. 14 (2023).

18. N. G. Kunigk, M. E. Urdaneta, I. G. Malone, F. Delgado, K. J. Otto, Reducing Behavioral Detection Thresholds per Electrode via Synchronous, Spatially-Dependent Intracortical Microstimulation. Front. Neurosci. 16, 876142 (2022).

19. M. E. Urdaneta, N. G. Kunigk, S. Currlin, F. Delgado, S. I. Fried, K. J. Otto, The Long-Term Stability of Intracortical Microstimulation and the Foreign Body Response Are Layer Dependent. Front. Neurosci. 16, 908858 (2022).

20. P. J. Rousche, R. A. Normann, Chronic intracortical microstimulation (ICMS) of cat sensory cortex using the Utah intracortical electrode array. IEEE Trans. Rehabil. Eng. 7, 56–68 (1999).

21. T. J. Smith, Y. Wu, C. Cheon, A. A. Khan, H. Srinivasan, J. R. Capadona, S. F. Cogan, J. J. Pancrazio, C. T. Engineer, A. G. Hernandez-Reynoso, Behavioral paradigm for the evaluation of stimulation-evoked somatosensory perception thresholds in rats. Serotonin Recept. Neurobiol. 17, 1202258 (2023).

22. R. Lycke, R. Kim, P. Zolotavin, J. Montes, Y. Sun, A. Koszeghy, E. Altun, B. Noble, R. Yin, F. He, N. Totah, C. Xie, L. Luan, Low-threshold, high-resolution, chronically stable intracortical microstimulation by ultraflexible electrodes. Cell Rep. 42, 112554 (2023).

23. T. Callier, E. W. Schluter, G. A. Tabot, L. E. Miller, F. V. Tenore, S. J. Bensmaia, Long-term stability of sensitivity to intracortical microstimulation of somatosensory cortex. J. Neural Eng. 12, 056010 (2015).

24. A. M. Ni, J. H. R. Maunsell, Microstimulation Reveals Limits in Detecting Different Signals from a Local Cortical Region. Curr. Biol. 20, 824–828 (2010).

25. J. R. Bartlett, E. A. DeYoe, R. W. Doty, B. B. Lee, J. D. Lewine, N. Negrão, W. H. Overman, Psychophysics of Electrical Stimulation of Striate Cortex in Macaques. J. Neurophysiol. 94, 3430–3442 (2005).

26. J. J. Cone, A. M. Ni, K. Ghose, J. H. R. Maunsell, Electrical Microstimulation of Visual Cerebral Cortex Elevates Psychophysical Detection Thresholds. eneuro 5, ENEURO.0311-18.2018 (2018).

27. A. S. Aberra, A. V. Peterchev, W. M. Grill, Biophysically realistic neuron models for simulation of cortical stimulation. J. Neural Eng. 15, 066023 (2018).

28. L. Sheintuch, A. Rubin, N. Brande-Eilat, N. Geva, N. Sadeh, O. Pinchasof, Y. Ziv, Tracking the Same Neurons across Multiple Days in Ca2+ Imaging Data. Cell Rep. 21, 1102–1115 (2017).

29. H. Zhu, F. He, P. Zolotavin, S. Patel, A. S. Tolias, L. Luan, C. Xie, Temporal coding carries more stable cortical visual representations than firing rate over time. Nat. Commun. 16, 7162 (2025).

30. L. Luan, J. T. Robinson, B. Aazhang, T. Chi, K. Yang, X. Li, H. Rathore, A. Singer, S. Yellapantula, Y. Fan, Z. Yu, C. Xie, Recent Advances in Electrical Neural Interface Engineering: Minimal Invasiveness, Longevity, and Scalability. Neuron 108, 302–321 (2020).

31. C. L. Hughes, K. A. Stieger, K. Chen, A. L. Vazquez, T. D. Kozai, Spatiotemporal properties of cortical excitatory and inhibitory neuron activation by sustained and bursting electrical microstimulation. iScience 28 (2025).

32. R. Yin, B. C. Noble, F. He, P. Zolotavin, H. Rathore, Y. Jin, N. Sevilla, C. Xie, L. Luan, Chronic co-implantation of ultraflexible neural electrodes and a cranial window. Neurophotonics 9 (2022).

33. F. He, Y. Sun, Y. Jin, R. Yin, H. Zhu, H. Rathore, C. Xie, L. Luan, Longitudinal neural and vascular recovery following ultraflexible neural electrode implantation in aged mice. Biomaterials 291, 121905 (2022).

34. L. Luan, X. Wei, Z. Zhao, J. J. Siegel, O. Potnis, C. A. Tuppen, S. Lin, S. Kazmi, R. A. Fowler, S. Holloway, A. K. Dunn, R. A. Chitwood, C. Xie, Ultraflexible nanoelectronic probes form reliable, glial scar–free neural integration. Sci. Adv. 3, e1601966 (2017).

35. Y. Wu, B. A. Temple, N. Sevilla, J. Zhang, H. Zhu, P. Zolotavin, Y. Jin, D. Duarte, E. Sanders, E. Azim, others, Ultraflexible electrodes for recording neural activity in the mouse spinal cord during motor behavior. Cell Rep. 43 (2024).

36. F. He, R. Lycke, M. Ganji, C. Xie, L. Luan, Ultraflexible Neural Electrodes for Long-Lasting Intracortical Recording. iScience 23, 101387 (2020).

37. C. Orlemann, C. Boehler, R. N. Kooijmans, B. Li, M. Asplund, P. R. Roelfsema, Flexible Polymer Electrodes for Stable Prosthetic Visual Perception in Mice. Adv. Healthc. Mater., 2304169 (2024).

38. R. Hattori, T. Komiyama, PatchWarp: Corrections of non-uniform image distortions in two-photon calcium imaging data by patchwork affine transformations. *Cell Rep*. Methods 2, 100205 (2022).

39. Q. Chen, J. Cichon, W. Wang, L. Qiu, S.-J. R. Lee, N. R. Campbell, N. DeStefino, M. J. Goard, Z. Fu, R. Yasuda, L. L. Looger, B. R. Arenkiel, W.-B. Gan, G. Feng, Imaging Neural Activity Using Thy1-GCaMP Transgenic Mice. Neuron 76, 297–308 (2012).

40. Y. Senzai, A. Fernandez-Ruiz, G. Buzsáki, Layer-Specific Physiological Features and Interlaminar Interactions in the Primary Visual Cortex of the Mouse. Neuron 101, 500–513.e5 (2019).

41. O. Kotler, Y. Khrapunsky, I. Fleidervish, Measuring Action Potential Propagation Velocity in Murine Cortical Axons. BIO-Protoc. 13 (2023).

42. M. C. Dadarlat, Y. J. Sun, M. P. Stryker, Activity-dependent recruitment of inhibition and excitation in the awake mammalian cortex during electrical stimulation. Neuron, S0896627323009273 (2023).

43. S. B. Hofer, H. Ko, B. Pichler, J. Vogelstein, H. Ros, H. Zeng, E. Lein, N. A. Lesica, T. D. Mrsic-Flogel, Differential connectivity and response dynamics of excitatory and inhibitory neurons in visual cortex. Nat. Neurosci. 14, 1045–1052 (2011).

44. E. J. Tehovnik, A. S. Tolias, F. Sultan, W. M. Slocum, N. K. Logothetis, Direct and Indirect Activation of Cortical Neurons by Electrical Microstimulation. J. Neurophysiol. 96, 512–521 (2006).

45. Y. Wang, Y. Chen, L. Chen, B. J. Herron, X. Y. Chen, J. R. Wolpaw, Motor learning changes the axon initial segment of the spinal motoneuron. J. Physiol. 602, 2107–2126 (2024).

46. S. Musall, M. T. Kaufman, A. L. Juavinett, S. Gluf, A. K. Churchland, Single-trial neural dynamics are dominated by richly varied movements. Nat. Neurosci. 22, 1677–1686 (2019).

47. C. Stringer, M. Pachitariu, N. Steinmetz, C. B. Reddy, M. Carandini, K. D. Harris, Spontaneous behaviors drive multidimensional, brainwide activity. Science 364, eaav7893 (2019).

48. E. H. Van Beest, C. Bimbard, J. M. J. Fabre, S. W. Dodgson, F. Takács, P. Coen, A. Lebedeva, K. D. Harris, M. Carandini, Tracking neurons across days with high-density probes. Nat. Methods 22, 778–787 (2025).

49. J. Shlens, Notes on Kullback-Leibler Divergence and Likelihood. arXiv arXiv:1404.2000 [Preprint] (2014). 10.48550/arXiv.1404.2000.

50. E. Maris, R. Oostenveld, Nonparametric statistical testing of EEG- and MEG-data. J. Neurosci. Methods 164, 177–190 (2007).

51. M. Okun, N. A. Steinmetz, L. Cossell, M. F. Iacaruso, H. Ko, P. Barthó, T. Moore, S. B. Hofer, T. D. Mrsic-Flogel, M. Carandini, K. D. Harris, Diverse coupling of neurons to populations in sensory cortex. Nature 521, 511–515 (2015).

52. S. N. Flesher, J. L. Collinger, S. T. Foldes, J. M. Weiss, J. E. Downey, E. C. Tyler-Kabara, S. J. Bensmaia, A. B. Schwartz, M. L. Boninger, R. A. Gaunt, Intracortical microstimulation of human somatosensory cortex. Sci. Transl. Med. 8 (2016).

53. M. Armenta Salas, L. Bashford, S. Kellis, M. Jafari, H. Jo, D. Kramer, K. Shanfield, K. Pejsa, B. Lee, C. Y. Liu, R. A. Andersen, Proprioceptive and cutaneous sensations in humans elicited by intracortical microstimulation. eLife 7, e32904 (2018).

54. E. Fernández, A. Alfaro, C. Soto-Sánchez, P. Gonzalez-Lopez, A. M. Lozano, S. Peña, M. D. Grima, A. Rodil, B. Gómez, X. Chen, P. R. Roelfsema, J. D. Rolston, T. S. Davis, R. A. Normann, Visual percepts evoked with an intracortical 96-channel microelectrode array inserted in human occipital cortex. J. Clin. Invest. 131, e151331 (2021).

55. G. Valle, A. H. Alamri, J. E. Downey, R. Lienkämper, P. M. Jordan, A. R. Sobinov, L. J. Endsley, D. Prasad, M. L. Boninger, J. L. Collinger, P. C. Warnke, N. G. Hatsopoulos, L. E. Miller, R. A. Gaunt, C. M. Greenspon, S. J. Bensmaia, Tactile edges and motion via patterned microstimulation of the human somatosensory cortex. Science 387, 315–322 (2025).

56. J. M. Rebesco, L. E. Miller, Enhanced detection threshold for *in vivo* cortical stimulation produced by Hebbian conditioning. J. Neural Eng. 8, 016011 (2011).

57. K. Ganguly, L. Kiss, M. Poo, Enhancement of presynaptic neuronal excitability by correlated presynaptic and postsynaptic spiking. Nat. Neurosci. 3, 1018–1026 (2000).

58. T. Alejandre-García, S. Kim, J. Pérez-Ortega, R. Yuste, Intrinsic excitability mechanisms of neuronal ensemble formation. eLife 11 (2022).

59. M. S. Grubb, Y. Shu, H. Kuba, M. N. Rasband, V. C. Wimmer, K. J. Bender, Short- and Long-Term Plasticity at the Axon Initial Segment. J. Neurosci. 31, 16049–16055 (2011).

60. A. T. Gulledge, J. J. Bravo, Neuron Morphology Influences Axon Initial Segment Plasticity. eneuro 3, ENEURO.0085-15.2016 (2016).

61. T. S. Davis, R. A. Parker, P. A. House, E. Bagley, S. Wendelken, R. A. Normann, B. Greger, Spatial and temporal characteristics of V1 microstimulation during chronic implantation of a microelectrode array in a behaving macaque. J. Neural Eng. 9, 065003 (2012).

62. S. P. Peron, J. Freeman, V. Iyer, C. Guo, K. Svoboda, A Cellular Resolution Map of Barrel Cortex Activity during Tactile Behavior. Neuron 86, 783–799 (2015).

63. J. Poort, A. G. Khan, M. Pachitariu, A. Nemri, I. Orsolic, J. Krupic, M. Bauza, M. Sahani, G. B. Keller, T. D. Mrsic-Flogel, S. B. Hofer, Learning Enhances Sensory and Multiple Non-sensory Representations in Primary Visual Cortex. Neuron 86, 1478–1490 (2015).

64. C. Verbaarschot, V. Karapetyan, C. M. Greenspon, M. L. Boninger, S. J. Bensmaia, B. Sorger, R. A. Gaunt, Conveying tactile object characteristics through customized intracortical microstimulation of the human somatosensory cortex. Nat. Commun. 16, 4017 (2025).

65. R. Kim, Y. Liu, J. Zhang, C. Xie, L. Luan, Towards precise synthetic neural codes: high-dimensional stimulation with flexible electrodes. *Npj Flex*. Electron. 9, 68 (2025).

66. A. Gilad, F. Helmchen, Spatiotemporal refinement of signal flow through association cortex during learning. Nat. Commun. 11, 1744 (2020).

67. The International Brain Laboratory, V. Aguillon-Rodriguez, D. Angelaki, H. Bayer, N. Bonacchi, M. Carandini, F. Cazettes, G. Chapuis, A. K. Churchland, Y. Dan, E. Dewitt, M. Faulkner, H. Forrest, L. Haetzel, M. Häusser, S. B. Hofer, F. Hu, A. Khanal, C. Krasniak, I. Laranjeira, Z. F. Mainen, G. Meijer, N. J. Miska, T. D. Mrsic-Flogel, M. Murakami, J.-P. Noel, A. Pan-Vazquez, C. Rossant, J. Sanders, K. Socha, R. Terry, A. E. Urai, H. Vergara, M. Wells, C. J. Wilson, I. B. Witten, L. E. Wool, A. M. Zador, Standardized and reproducible measurement of decision-making in mice. eLife 10, e63711 (2021).

68. T. Zito, D.-E. Künstle, G. Aguilar, P. Berkes, L. Schwetlick, psignifit 4.3, version v4.3, Zenodo (2025); 10.5281/ZENODO.14765353.

69. H. H. Schütt, S. Harmeling, J. H. Macke, F. A. Wichmann, Painfree and accurate Bayesian estimation of psychometric functions for (potentially) overdispersed data. Vision Res. 122, 105–123 (2016).

70. F. Dinc, H. Inan, O. Hernandez, C. Schmuckermair, O. Hazon, T. Tasci, B. O. Ahanonu, Y. Zhang, J. Lecoq, S. Haziza, M. J. Wagner, M. A. Erdogdu, M. J. Schnitzer, Fast, scalable, and statistically robust cell extraction from large-scale neural calcium imaging datasets. Neuroscience [Preprint] (2021). 10.1101/2021.03.24.436279.

71. H. Inan, M. A. Erdogdu, M. Schnitzer, “Robust Estimation of Neural Signals in Calcium Imaging” in Advances in Neural Information Processing Systems (NeurIPS 2017) (2017)vol. 30.

72. J. Friedrich, P. Zhou, L. Paninski, Fast online deconvolution of calcium imaging data. PLOS Comput. Biol. 13, e1005423 (2017).

