## Supplementary Material for "Learning induces activation-mechanism–dependent neural plasticity in an intracortical microstimulation task"

Science Advances

AAAS

Robin Kim *et al.*

**This PDF file includes:**

Figs. S1 to S9

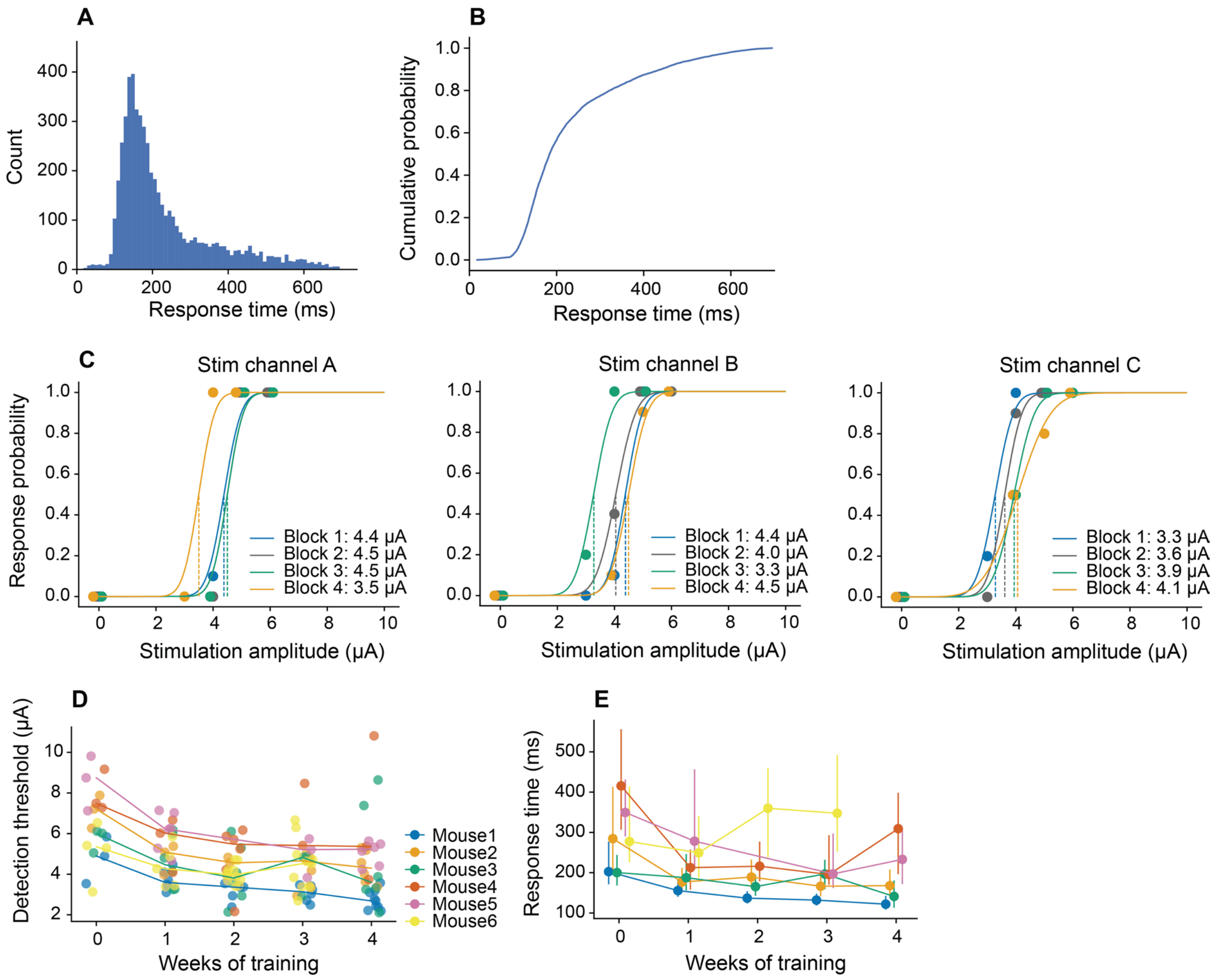


**Fig. S1. Behavioral metrics of trained mice.** (**A**) Distribution of response times for all hit trials during detection threshold measurements. (**B**) Cumulative distribution function (CDF) of response times shown in (**A**). (**C**) Psychometric curves from three stimulation channels across four blocks in an example session. Within-session threshold drifts motivated dividing the session into four blocks; stimulation amplitudes were adjusted based on the response probabilities in the previous block to maintain peri-threshold stimulation. (**D**) Detection thresholds for each stimulation channel across all animals and sessions. Each point represents a stimulation channel threshold for a single session; lines denote median values. (**E**) Response times across training for individual animals. Points represent medians; error bars indicate interquartile ranges (IQRs). n = 6 animals for 46 sessions.

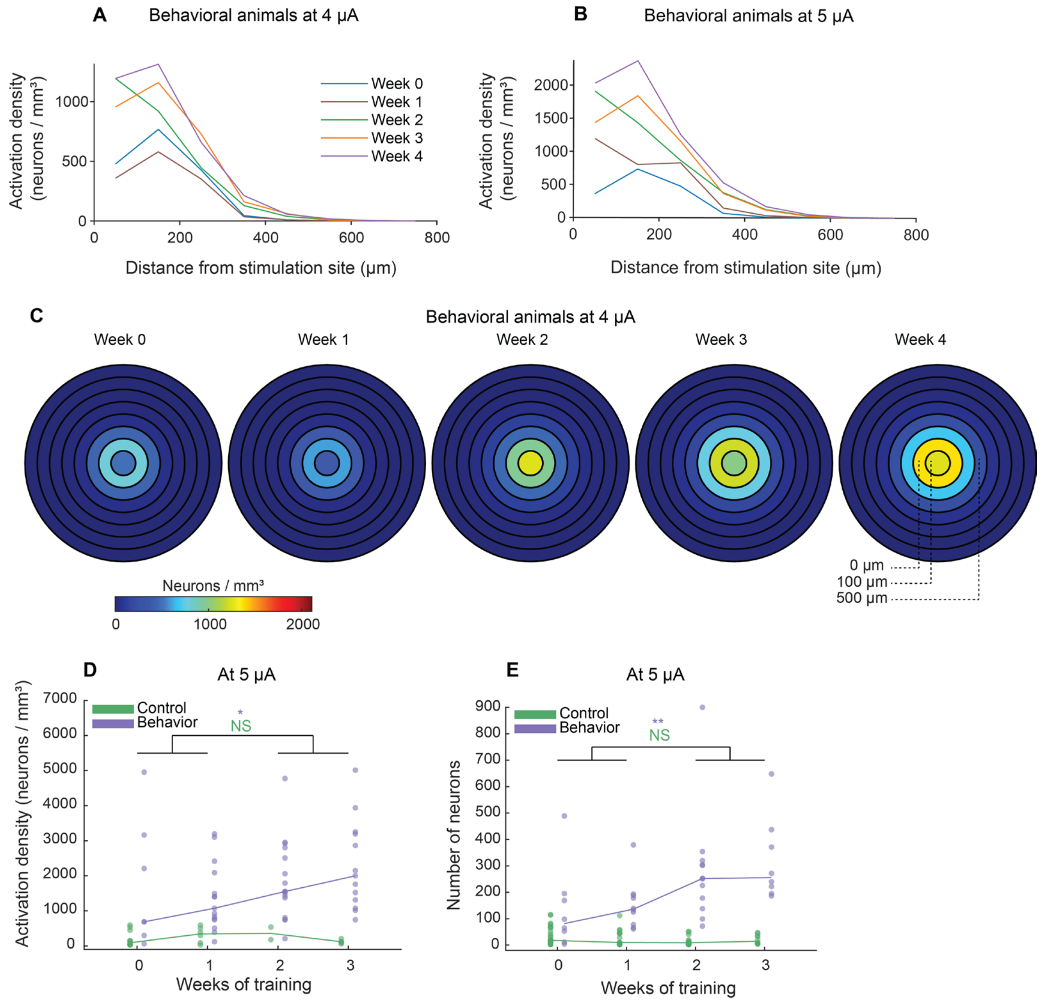


**Fig. S2. Neural activation and spatial spread at other currents and in control animals.** (**A, B**) Activation density as a function of distance to the stimulation sites across training periods for 4 and 5 μA, respectively. (**C**) Ring plots at 4 μA showing spatial distribution of ICMS-evoked activation. Recruitment increases over weeks of behavioral learning, consistent with the results at 5 μA shown in Fig. 2D. (**D**) In non-behavioral control animals, activation density within 200 μm of the stimulation site remained stable over weeks of stimulation (two-sided Mann-Whitney U, $n_{early} = 18$, $n_{late} = 7$, $p = 0.808$, $r = 0.05$), whereas behaviorally trained animals exhibited significant increases in activation density (two-sided Mann-Whitney U, $n_{early} = 22$, $n_{late} = 29$, $p = 0.044$, $r = 0.28$). Activation density data from behaviorally trained animals are reproduced from Fig. 2E for comparison. (**E**) Similarly, non-behavioral controls recruited a stable population over weeks of stimulation (two-sided Mann-Whitney U; $n_{early} = 42$, $n_{late} = 26$, $p = 0.066$, $r = 0.22$), while behaviorally trained animals showed significant increases in overall population recruitment at 5 μA (two-sided Mann-Whitney U, $n_{early} = 18$, $n_{late} = 20$, $p = 9.0\text{e-}4$, $r = 0.54$). Neuron count data from behaviorally trained animals are reproduced from Fig. 2C for comparison. n = 6 animals for 46 sessions; n = 6 control animals for 18 sessions.

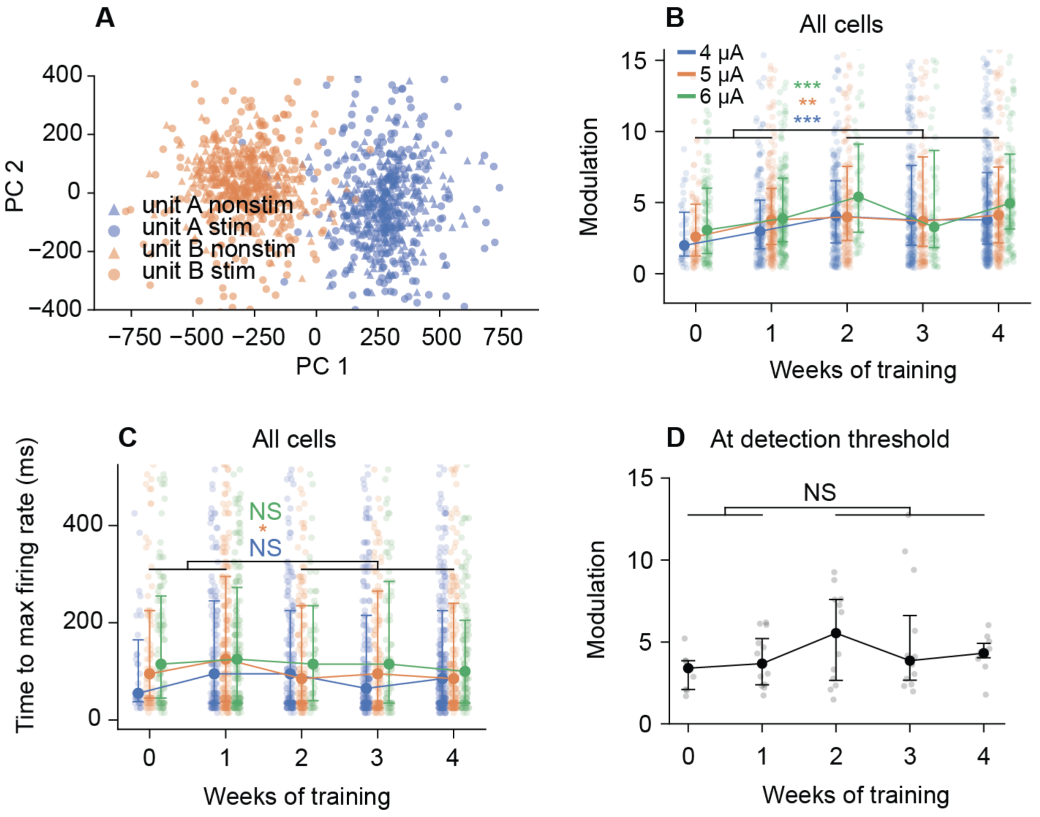


**Fig. S3. Spike sorting validation and change of modulation over learning of all modulated units.** (**A**) PCA of spike waveforms from two example units, subdivided into spikes recorded during stimulation and non-stimulation periods. For each unit, stim and non-stim spikes cluster together, confirming waveform stability across conditions. (**B**) Modulation z-score at fixed currents of all modulated cells increased significantly over training (two-sided Mann-Whitney U; 4 µA: $n_{early} = 189$, $n_{late} = 729$, $p = 4.8e\text{-}5$, $r = 0.18$; 5 µA: $n_{early} = 283$, $n_{late} = 489$, $p = 3.3e\text{-}3$, $r = 0.12$; 6 µA: $n_{early} = 303$, $n_{late} = 250$, $p = 1.5e\text{-}4$, $r = 0.18$). (**C**) Time to maximum firing rate of all modulated units. A decrease was observed at 5 µA ($p = 0.012$, $r = -0.10$) but not with statistical significance at 4 µA ($p = 0.18$, $r = -0.04$) or 6 µA ($p = 0.19$, $r = -0.04$). (**D**) Modulation at each session's psychometric detection threshold remained stable over training (two-sided Mann-Whitney U, $n_{early} = 21$, $n_{late} = 34$, $p = 0.095$, $r = 0.21$). Medians ± IQR are shown throughout; individual observations are plotted with jitter. n = 6 animals for 46 sessions.

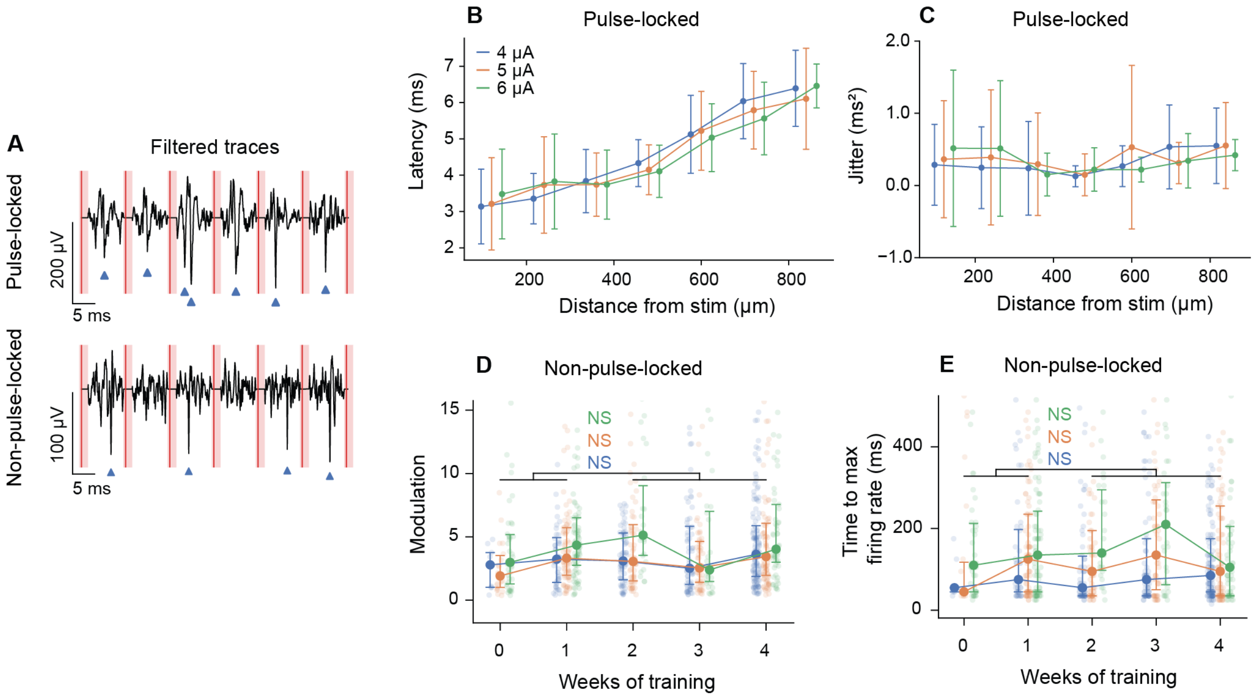


**Fig. S4. Pulse-locked and non-pulse-locked response metrics.** (**A**) Preprocessed voltage traces (see Fig. 5A for raw traces) from a representative pulse-locked (PL, top) and non-pulse-locked (NPL, bottom) unit at 5 μA. Detected spikes (blue triangles) are aligned with threshold-crossing events of similar amplitude, validating spike detection for both PL and NPL cells. Red vertical lines indicate ICMS pulses; light red shading denotes blanking periods. (**B**) Latency of PL responses as a function of distance from the stimulation site, separated by current amplitude. OLS regression on binned means (120 μm bins) showed a significant linear increase at all currents (4 μA: slope = 0.005 ms/μm, $p < 0.001$, $r^2 = 0.97$, speed ~ 0.20 m/s; 5 μA: slope = 0.004 ms/μm, $p < 0.001$, $r^2 = 0.95$, speed ~ 0.24 m/s; 6 μA: slope = 0.004 ms/μm, $p = 0.001$, $r^2 = 0.91$, speed ~ 0.25 m/s). (**C**) Jitter of PL responses as a function of distance from the stimulation site. OLS regression on binned means showed no significant relationship at any current level (4 μA: $p = 0.095$; 5 μA: $p = 0.43$; 6 μA: $p = 0.53$). (**D**) NPL modulation z-score over weeks of training. No significant increase was observed at any current level (two-sided Mann-Whitney U; 4 μA: $n_{early} = 53$, $n_{late} = 260$, $p = 0.093$, $r = 0.12$; 5 μA: $n_{early} = 95$, $n_{late} = 204$, $p = 0.38$, $r = 0.02$; 6 μA: $n_{early} = 124$, $n_{late} = 115$, $p = 0.15$, $r = 0.08$). (**E**) NPL time to maximum firing rate over weeks. No significant change was observed (two-sided Mann-Whitney U; 4 μA: $p = 0.38$, $r = -0.03$; 5 μA: $p = 0.51$, $r = 0.00$; 6 μA: $p = 0.85$, $r = 0.08$). Medians ± IQR are shown throughout; individual observations are plotted with jitter. n = 6 animals for 46 sessions.

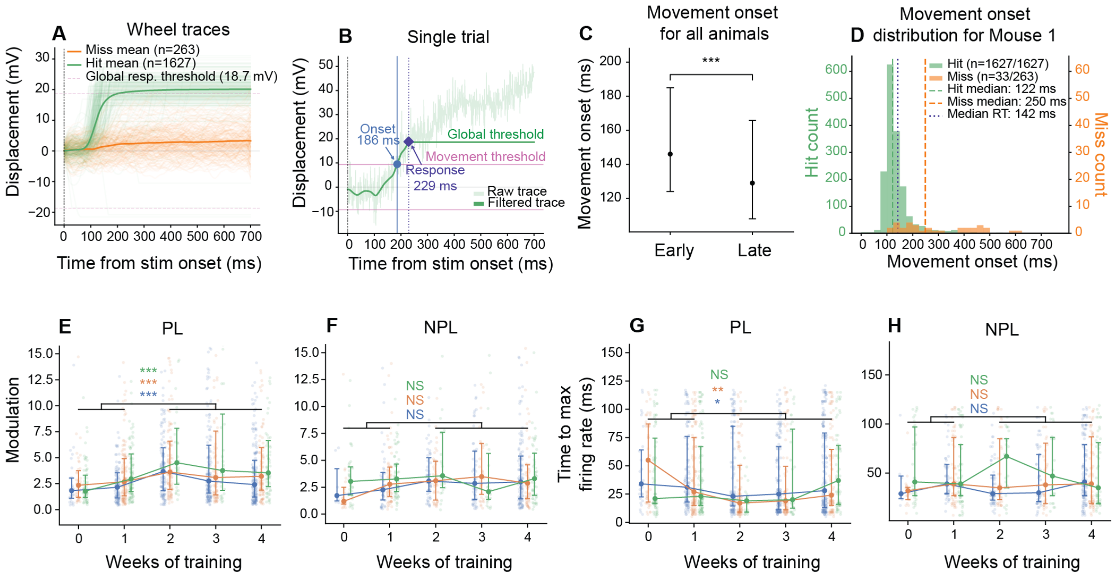


**Fig. S5. Neural modulation in PL and NPL cells in a restricted time window designed to minimize movement-related confounds.** (**A**) Wheel displacement traces for all hit (green; n = 1,627) and miss (orange; n = 263) trials from Mouse 1, which had the fastest response time among all trained animals (4-6 µA, 10 sessions). Dashed lines indicate the global response threshold (25th percentile of absolute displacement at response time). (**B**) Representative single trial illustrating the movement detection method. A 20 Hz low-pass forward-backward filtered displacement trace (solid) is shown alongside the raw trace (transparent). Movement onset (blue dot and vertical line) is defined as the first time point at which displacement exceeds 50% of the global response threshold. The behavioral response time is marked by the purple diamond. (**C**) Wheel-movement onset times decrease from early to late training (two-sided Mann-Whitney U; $n_{early}$ =1,305 trials, $n_{late}$ = 2,922 trials, p = 4.15e-30, r = -0.22). Medians ± IQR are shown. n = 5 animals, 38 sessions; Mouse 6 was excluded due to reduced encoder dynamic range. (**D**) Distribution of movement onset latencies for hit and miss trials for Mouse 1. Median hit onset (122 ms) and median response time of all trials (142 ms) are indicated by dashed and dotted vertical lines, respectively. (**E**) PL modulation z-score across weeks of training, computed using the restricted window. PL modulation increased significantly at all current levels (two-sided Mann-Whitney U; 4 µA: $n_{early}$ = 127, $n_{late}$ = 435, p = 2.2e-5, r = 0.25; 5 µA: $n_{early}$ = 164, $n_{late}$ = 276, p = 4.7e-4, r = 0.20; 6 µA: $n_{early}$ = 165, $n_{late}$ = 127, p = 2.0e-5, r = 0.29). (**F**) NPL modulation z-score. NPL modulation did not show a significant change at any current level (4 µA: $n_{early}$ = 52, $n_{late}$ = 225, p = 0.075, r = 0.16; 5 µA: $n_{early}$ = 83, $n_{late}$ = 147, p = 0.089, r = 0.14; 6 µA: $n_{early}$ = 103, $n_{late}$ = 104, p = 0.81, r = 0.02). (**G**) PL time to maximum firing rate. Time to maximum decreased at 4 µA (p = 0.038, r = -0.12) and 5 µA (p = 1.7e-3, r = -0.18) but not 6 µA (p = 0.77, r = -0.02). (**H**) NPL time to maximum firing rate, which showed no significant change at any current level (4 µA: p = 0.45, r = -0.07; 5 µA: p = 0.90, r = -0.01; 6 µA: p = 0.70, r = -0.03). Medians ± IQR are shown throughout; individual observations are plotted with jitter. n = 6 animals, 46 sessions for **E-H**.

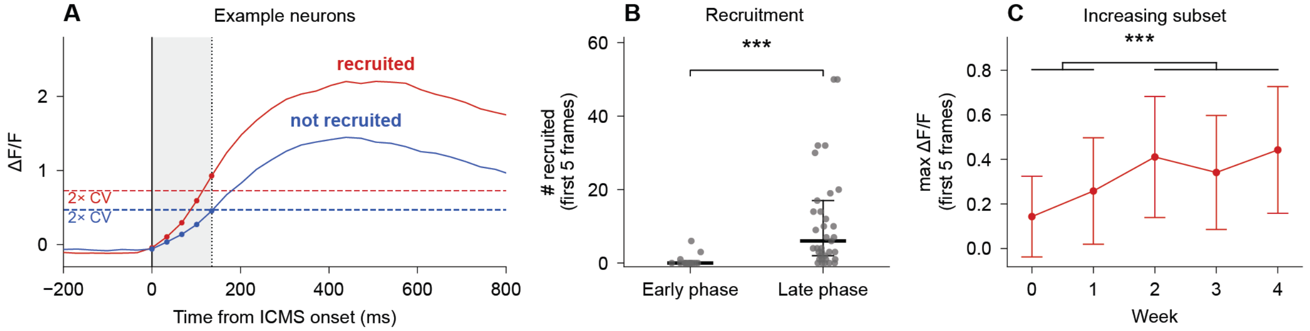


**Fig. S6. Two-photon imaging shows an increase in neuronal recruitment and response amplitude with training within the first five frames following ICMS onset.** (**A**) Two example trial-averaged ΔF/F traces from individual neurons: one whose response crosses threshold within the first five frames (red) and one that does not (blue). A neuron is considered recruited at a given stimulation channel × current if its trial-averaged ΔF/F exceeds 2 × CV in any of the first five frames where CV is the coefficient of variation (standard deviation / mean) of the trial-concatenated fluorescence trace. (**B**) The number of recruited neurons per stimulation channel × session increased significantly from early to late training (two-sided Mann-Whitney U; nearly = 15, nlate = 37, p = 1.8e-5, r = 0.75). Each dot represents one (animal × session × stimulation channel); value is the number of tracked neurons recruited at 4-6 µA. Medians ± IQR are shown. (**C**) Maximum ΔF/F within the first five frames for the increasing subset defined in Fig. 3J, across weeks of training (two-sided Mann-Whitney U; p = 4.1e-4, r = 0.63). Means ± SD are shown.

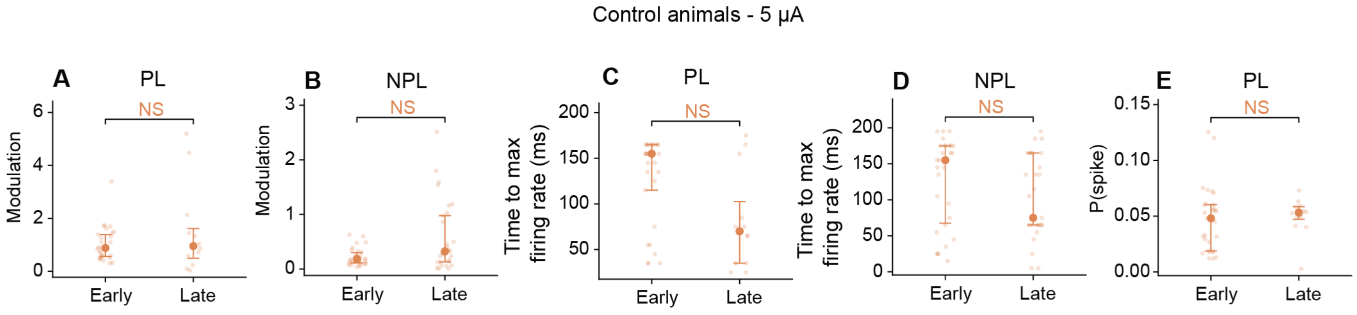


**Fig. S7. ICMS in the control group (without behavioral training) does not induce significant changes in neural modulation.** (**A**) PL modulation z-score ($n_{early}$ = 29, $n_{late}$ = 12, p = 0.83, r = 0.05). (**B**) NPL modulation z-score ($n_{early}$ = 30, $n_{late}$ = 29, p = 0.061, r = 0.29). **(C)** PL time to maximum firing rate (p = 0.050, r = -0.39). (**D**) NPL time to maximum firing rate (p = 0.13, r = -0.23). (**E**) PL maximum spike probability (p = 0.62, r = 0.10). All comparisons use two-sided Mann-Whitney U tests. Medians ± IQR are shown; individual observations are plotted with jitter. n = 4 control animals, 15 sessions.

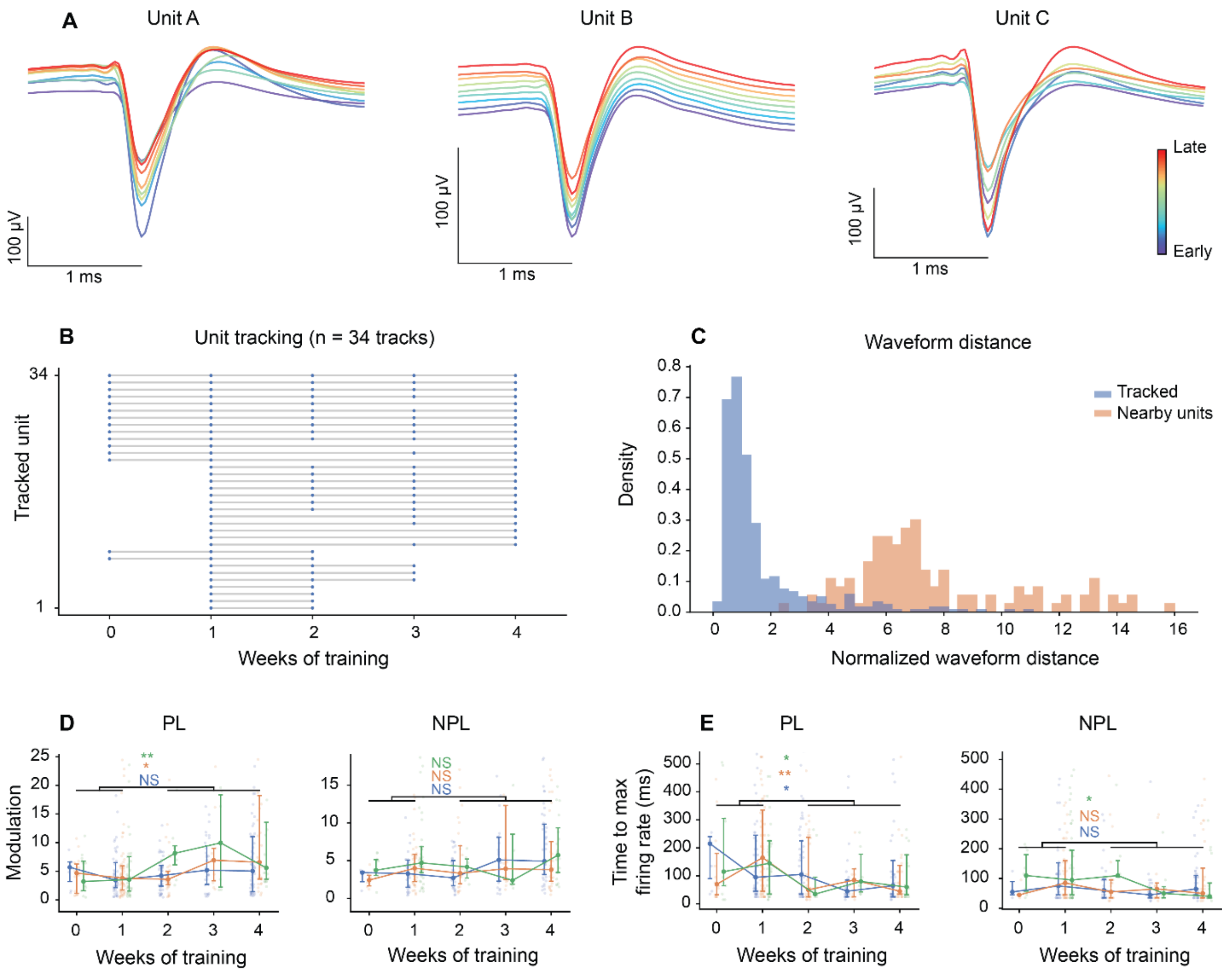


**Fig. S8. Electrophysiological unit tracking.** (**A**) Example waveforms from three tracked units recorded across sessions. Colors indicate recording session (early to late). (**B**) Raster of all tracked units. Each row represents a single track. Each dot marks week in which the unit was detected. Units that appeared in both early (weeks 0-1) and late (weeks 2-4) learning phases are included. (**C**) Distribution of normalized waveform distances for tracked pairs (within-unit, cross-session) and nearby non-matched units (different units detected by nearby channels, same session). Tracked pairs were 6.8-fold more similar than nearby non-matched units (two-sided Mann-Whitney U; $p = 6.6e-83$), confirming reliable unit identification. (**D**) Longitudinal modulation changes for tracked units. PL modulation z-score increased significantly from early to late learning phases (two-sided Mann-Whitney U; 5 µA: $p = 0.049$, $r = 0.19$; 6 µA: $p = 0.0024$, $r = 0.39$), while NPL modulation did not show significant changes. PL time to maximum firing rate decreased significantly at all current levels (4 µA: $p = 0.026$, $r = -0.21$; 5 µA: $p = 0.0028$, $r = -0.28$; 6 µA: $p = 0.030$, $r = -0.28$). Medians ± IQR are shown throughout; individual observations are plotted with jitter. n = 34 tracked units, 6 animals, 46 sessions.

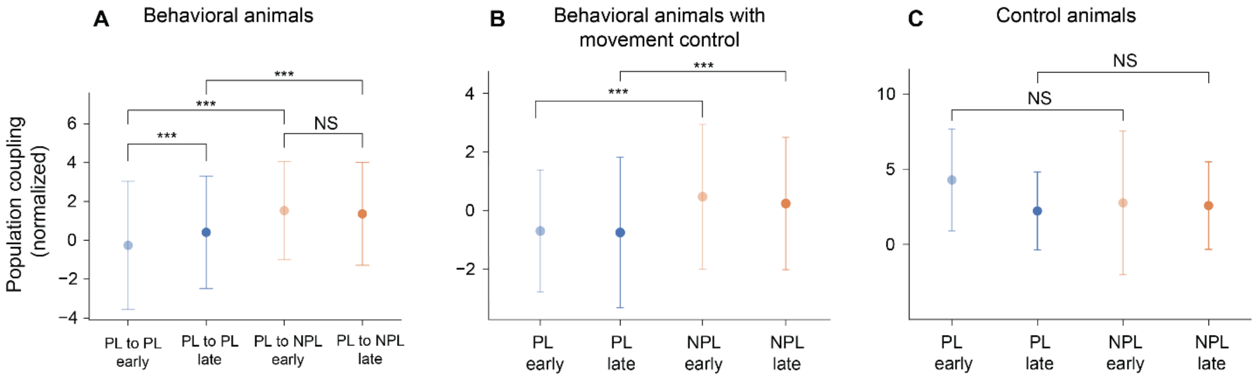


**Fig. S9. Population coupling analysis across experimental conditions.** (**A**) Population coupling (normalized) for PL and NPL units in early and late training phases (700 ms analysis window). PL coupling increased significantly over time (PL early vs late: $n_{early}$ = 446, $n_{late}$ = 763, $p_{FDR}$ = 1.3e-6, r = 0.17), while NPL coupling did not change (NPL early vs late: $n_{early}$ = 255, $n_{late}$ = 547, $p_{FDR}$ = 0.22, r = -0.05). PL coupling was significantly lower than NPL coupling in both early ($p_{FDR}$ = 5.5e-24, r = 0.46) and late ($p_{FDR}$ = 3.4e-17, r = 0.28) periods (two-sided Mann-Whitney U with FDR-BH correction). (**B**) Same analysis using the restricted 120 ms analysis window. PL coupling was significantly lower than NPL coupling in both early ($n_{PL}$ = 264, $n_{NPL}$ = 125, $p_{FDR}$ = 7.4e-9, r = 0.37) and late ($n_{PL}$ = 484, $n_{NPL}$ = 255, $p_{FDR}$ = 1.7e-11, r = 0.31) periods. **(C)** Population coupling in control animals (5 µA, no behavioral training). PL and NPL coupling did not differ significantly in either early ($n_{PL}$ = 49, $n_{NPL}$ = 64, $p_{FDR}$ = 0.48, r = -0.10) or late ($n_{PL}$ = 25, $n_{NPL}$ = 128, $p_{FDR}$ = 0.48, r = 0.13) periods. All comparisons use two-sided Mann-Whitney U with Benjamini-Hochberg correction. Medians ± IQR are shown. n = 6 animals, 46 sessions; n = 4 control animals, 15 sessions.
